# NUDT5 regulates the global efficacy of nucleoside analog drugs by coordinating purine synthesis and PRPP allocation

**DOI:** 10.1101/2025.11.20.689348

**Authors:** Nicholas C.K. Valerie, Seher Alam, Femke M. Hormann, Ulf Martens, Bo Lundgren, Massimiliano Gaetani, Johan Boström, Sean G. Rudd, Per I. Arvidsson, Mikael Altun

**Author notes:** Corresponding authors:* Nicholas C.K. Valerie and Mikael Altun; Karolinska Institutet, ANA Futura, Alfred Nobels Allé 8, 141 52 Huddinge, Sweden.

## Abstract

Cancer cells require nucleotides to meet proliferation demands, which has been effectively exploited by nucleoside analog (NA) drugs. Despite their success, we still do not completely understand factors dictating their anticancer effect. Here, we report that a NUDT5 scaffolding function indirectly regulates the global efficacy of anti-cancer NA drugs. Mechanistically, PROTAC- or RNAi-mediated NUDT5 depletion, not inhibition, desensitizes cells to 6-thioguanine (6TG) and other NA drugs proportionally with NUDT5 abundance. NUDT5-depleted cells appear locked into *de novo* purine biosynthesis (DNPB), thus impairing salvage of nucleotide precursors and NA drug activation. Specifically, NUDT5 interacts with phosphoribosyl amidotransferase (PPAT), the rate-limiting DNPB enzyme, to putatively regulate its activity, DNPB flux, and resultant phosphoribosyl pyrophosphate (PRPP) allocation. Collectively, these results suggest that NUDT5 controls NA drug efficacy by coordinating nucleotide synthesis and PRPP utilization, making it a potential biomarker of clinical efficacy and target for finetuning antimetabolite therapies.

**Statement of significance:** NUDT5 non-enzymatically regulates the global efficacy of anti-cancer nucleoside analog drugs by interacting with PPAT to control DNPB and allocation of PRPP, thereby establishing it as a biomarker of clinical significance.

## Introduction

The synthesis of deoxyribonucleotide triphosphates (dNTPs) is essential for DNA replication and repair and occurs through two interconnected routes: the *de novo* pathway, which constructs nucleotides from small molecule precursors, and the salvage pathway, which recycles nucleosides and nucleobases (1,2). For example, the *de novo* purine biosynthesis (DNPB) pathway generates inosine monophosphate (IMP) from phosphoribosyl pyrophosphate (PRPP) in a multi-step process starting with the rate-limiting enzyme, phosphoribosyl pyrophosphate amidotransferase (PPAT), followed by infusion of one-carbon units from the folate cycle (1). This energetically costly process requires ATP at several steps; thus, *de novo* and salvage pathways are dynamically tuned to the cellular metabolic state. Many cancers display increased glycolytic flux and metabolic reprogramming (*i.e.*, the Warburg effect) that shift this balance toward *de novo* synthesis to meet the high nucleotide demand of rapid cell division (3,4).

Despite differences in energetic requirements, pentose units are utilized by both pathways in the form of ribose-5-phosphate (R5P), synthesized primarily *via* the pentose phosphate pathway (PPP) (1). The oxidative branch of the PPP provides NADPH for reductive biosynthesis, while the non-oxidative branch interconverts glycolytic intermediates to supply R5P for PRPP synthesis under variable nutrient conditions. As PRPP is a shared substrate, its cellular pool is tightly regulated and fluctuations in its availability can shift the balance between the two routes (5–11).

Exploiting the dependency on active nucleotide metabolism has long been an attractive and effective anticancer strategy (2). For example, the *de novo* pathway has been targeted by antifolates, such as methotrexate (MTX), to deprive cancer cells of dNTPs. Conversely, the salvage pathway has been utilized for nucleoside analog (NA) drugs, which are prodrugs that must be phosphorylated to active metabolites to exert their effects (12). For example, thiopurines (6-thioguanine [6TG] and 6-mercaptopurine [6MP]) are mainstay therapeutics for childhood leukemia typically used in combination with *de novo* synthesis inhibitors (13). Despite their success, the determinants of NA drug efficacy remain incompletely understood – as evidenced by the ongoing discovery of NA drug metabolizers, such as SAMHD1 (14–16) and NUDT15 (17–20).

Recently, genome-wide loss-of-function screens have identified Nudix hydrolase 5 (NUDT5) as a potential enabler of thiopurine efficacy (21,22). NUDT5 hydrolyzes ADP-ribose (ADPr) to AMP and R5P but can also synthesize ATP from ADPr to fuel DNA repair and hormone-dependent transcription (23–27). Thus, it was inferred that NUDT5 ablation could blunt 6TG efficacy by limiting R5P required for HPRT-mediated activation, but this was never substantiated. Here, we report that a NUDT5 scaffolding function indirectly mediates the global anticancer efficacy of NA drugs by regulating DNPB activity, thereby enabling their salvage-dependent activation. This effect is titratable and directly correlated with NUDT5 protein abundance, suggesting NUDT5 may be a valuable prognostic biomarker of NA drug clinical efficacy and target for finetuning antimetabolite therapies.

## Results

### NUDT5 depletion, not inhibition, elicits thioguanine resistance

In addition to its reported roles in breast cancer cell transcriptional regulation (25,26) and DNA repair (27), NUDT5 has also appeared as a hit in numerous loss-of-function positive selection screens for 6-thioguanine (6TG) resistance (21,22). These studies proposed a mechanism related to NUDT5 catalysis, where NUDT5 loss-of-function could limit HPRT-mediated activation of 6TG (**Fig. 1A**) – nonetheless, this mechanism was never validated.

**Figure 1.**
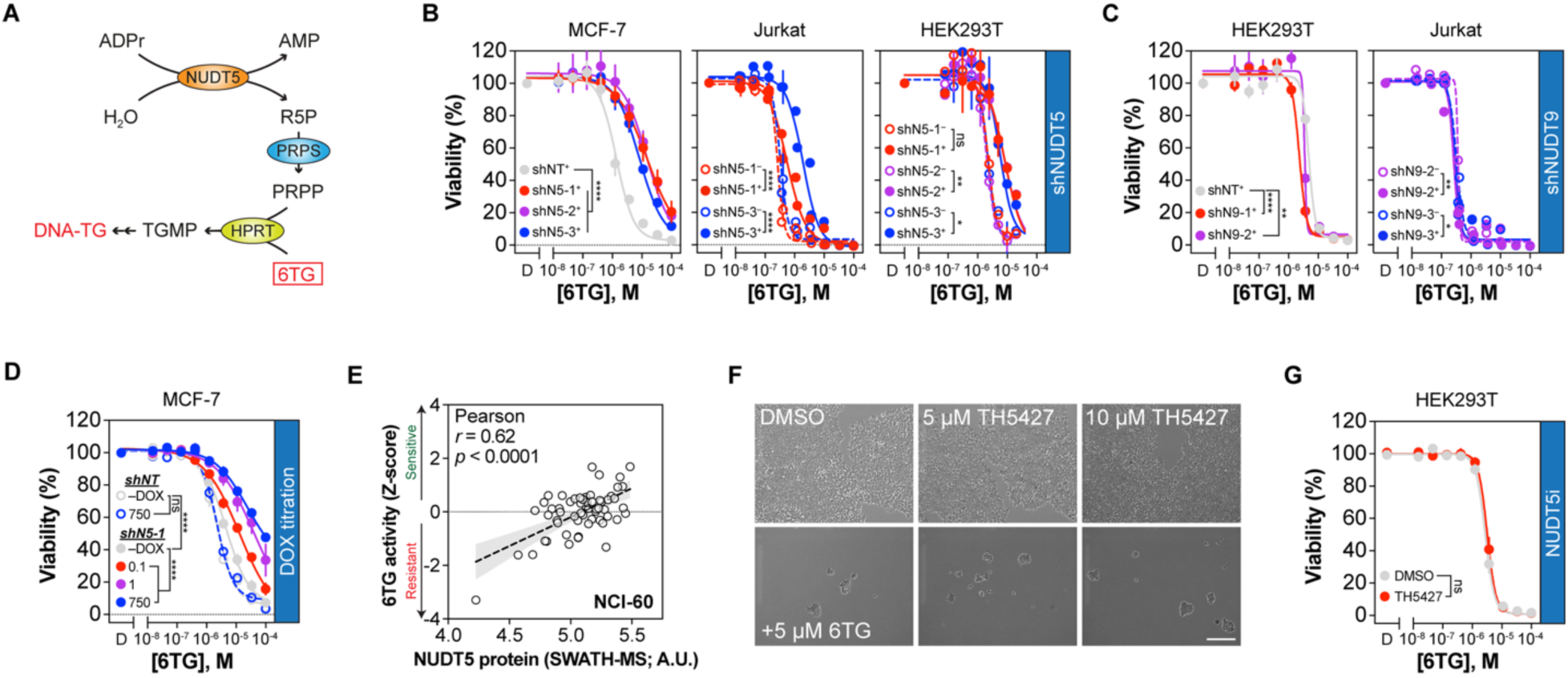
NUDT5 depletion, not inhibition, yields thioguanine resistance in cells. **A**, Proposed model for NUDT5 involvement in thioguanine (6TG) toxicity by Doench *et al.* and Sagi *et al.* (21,22). **B**, NUDT5 was depleted by doxycycline (DOX)-inducible shRNA (+; filled circles) in MCF-7 (n=3 repeats), Jurkat (n=2 repeats), or HEK293T (n=2 repeats) cells and incubated with a 6TG concentration gradient for 96 hours before viability measurement. No DOX (–) or non-targeting hairpins (shNT) were used as controls. **C**, NUDT9-depleted HEK293T or Jurkat cells (n=2 repeats) were incubated with a 6TG concentration gradient for 96 hours before viability measurement. **D**, Control or NUDT5 shRNA was induced in MCF-7 cells by DOX titration before incubation with a 6TG concentration gradient for 96 hours and viability measurement. **E**, Pearson correlation of NUDT5 protein expression (SWATH-MS) and 6TG activity in the NCI-60 dataset (*r* = 0.62, *p* < 0.001; linear regression ± 95% CI shown). **F**, Representative light contrast micrographs of HEK293T cells incubated with DMSO or TH5427 (5 or 10 µmol/L) ± 5 µmol/L 6TG for 72 hours. Scale bar = 200 µm. **G**, HEK293T cells were preincubated with DMSO or TH5427 (5 µmol/L) before adding a 6TG concentration gradient for 96 hours and viability measurement (means ± SEM from n=2 repeats). In all cases, statistical significance determined by extra sum-of-squares F-test for EC_50_ difference from control (**B**-**D**, **G**) or two-sided t test (**E**), and error reported as ± SEM.

We first sought to confirm loss-of-function screening data and performed 6TG dose-response viability analyses using NUDT5-targeted shRNA with doxycycline-regulable expression. Efficient knockdown by up to three different NUDT5 shRNAs was confirmed by both qPCR and protein expression in several cell lines (**Supplementary Fig. S1**). NUDT5 depletion significantly desensitized cells to 6TG, irrespective of viability changes from the individual shRNAs alone (**Fig. 1B, Supplementary Fig. S1H-K**). Meanwhile, depletion of NUDT9, a NUDT5 paralog also catalyzing ADP-ribose hydrolysis, had no discernible effect on 6TG toxicity, suggesting the effect is specific to NUDT5 and not disruption of ADP-ribose hydrolysis *per se* (**Fig. 1C, Supplementary Fig. S1B, H, and I**). Furthermore, 6TG resistance was titratable with increasing doxycycline concentrations, implying that NUDT5 abundance directly correlates with 6TG efficacy (**Fig. 1D**).

We then sought curated evidence that NUDT5 abundance could dictate the efficacy of 6TG and turned to the CellMinerCDB database, which curates molecular and pharmacological datasets for analysis (28). NUDT5 protein abundance positively correlated with 6TG (*r* = 0.62, *p* < 0.0001 [Pearson]; **Fig. 1E**) and 6-mercaptopurine (6MP; *r* = 0.36, *p* = 0.0056 [Pearson]; **Supplementary Fig. S2A**) activity but worse so by mRNA expression in the NCI-60 dataset and the larger CCLE/PRISM datasets (**Supplementary Fig. S2B-E**). We then tested if a previously reported NUDT5 inhibitor (TH5427) could also elicit 6TG resistance in cells (26). However, despite earlier confirmation that it potently engages NUDT5 in HEK293T cells (29), TH5427 had no effect on 6TG toxicity up to tested concentrations of 10 µmol/L (**Fig. 1F and G**). Thus, while NUDT5 abundance positively correlates with 6TG sensitivity, potent inhibitors are unable to recapitulate the elicited effect.

### NUDT5 PROTAC, DDD2, confirms that NUDT5 abundance, not activity, dictates 6TG toxicity

As NUDT5 inhibition failed to recreate the desensitization seen by RNAi, we sought an alternative pharmacological approach to target NUDT5 activities. Proteolysis targeting chimeras (PROTACs) affect all potential functions of the desired protein by routing it for destruction *via* targeted protein degradation (TPD). We recently identified a selective, VHL-directed PROTAC towards NUDT5 (DDD2) with a DC_50_ and D_max_ of up to ∼10 nM and ∼90%, respectively, using tunable degradation assays and a cell-first screening approach (**Fig. 2A**) (29). We then compared the effect of both NUDT5 targeting modalities on 6TG-mediated toxicity. As before, DDD2 durably degraded endogenous NUDT5 in multiple cell lines for up to several days, and, like with RNAi, resulted in significant (∼ 4 to 30-fold) resistance to 6TG, whereas TH5427 had no effect (**Fig. 2B-D**, **Supplementary Fig. S3A and B**). Notably, DDD2 or TH5427 alone had no obvious effects on cell viability at tested concentrations (**Supplementary Fig. S3C-E**). Similarly, DDD2-mediated 6TG resistance was titratable with a plateau at ∼5 µmol/L, coinciding with the reported NUDT5 degradation profile (**Fig. 2E**) (29). Importantly, we also saw that DDD2-dependent 6TG resistance could be reverted upon catalytic site competition with TH5427 (**Fig. 2F**), in line with TH5427 preventing DDD2-mediated NUDT5 degradation (29), which confirmed the effect is contingent on NUDT5 presence, not inhibition.

**Figure 2.**
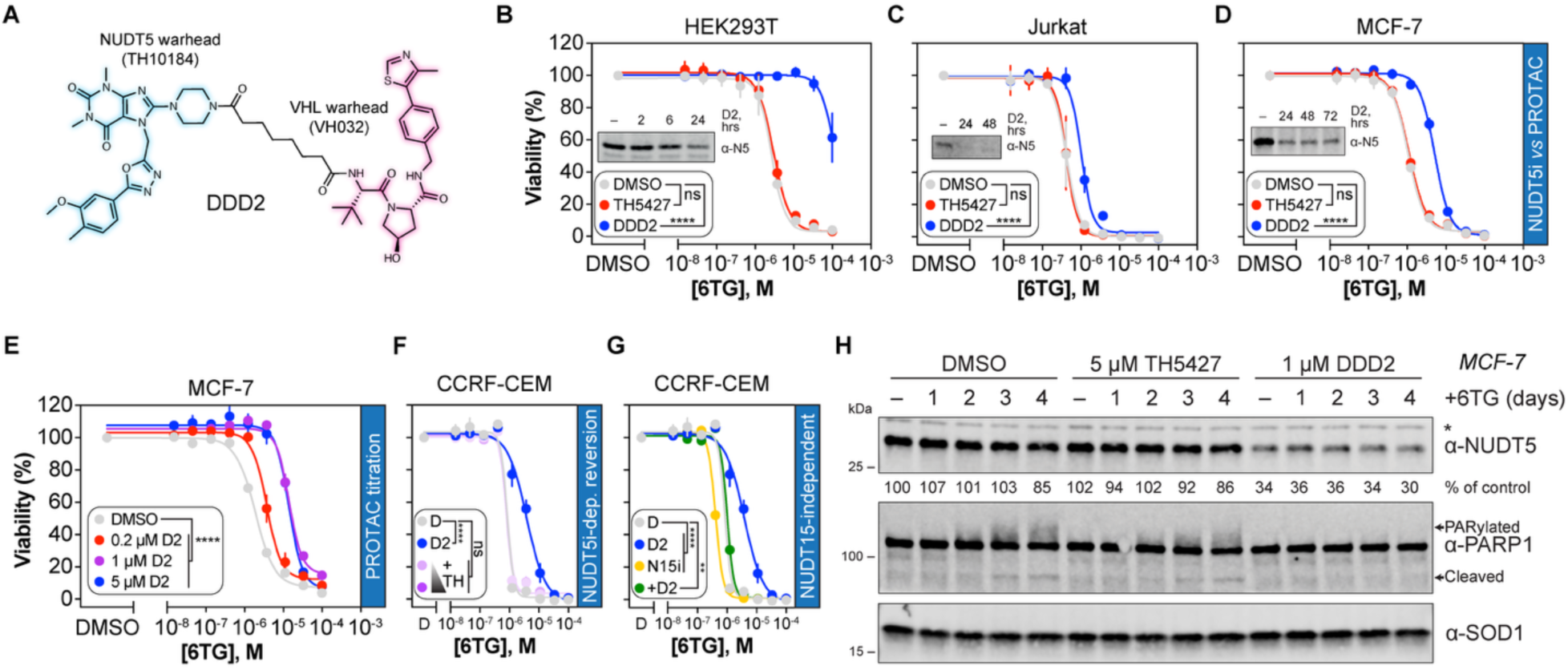
NUDT5 PROTAC, DDD2, phenocopies RNAi-mediated 6TG resistance in a NUDT5-dependent manner. **A**, Structure of NUDT5 PROTAC, DDD2, with NUDT5 (blue) and VHL warheads (magenta) highlighted. **B**, HEK293T (n=3 repeats), Jurkat (**C**, n=2 repeats), or MCF-7 cells (**D**, n=2 repeats) were pretreated with DMSO, TH5427 (5 µmol/L), or DDD2 (1 µmol/L) for 24 hours followed by a 6TG gradient for an additional 96 hours prior to viability measurement. Western blot insets depict confirmation of DDD2-mediated NUDT5 depletion (representative of n=2 repeats). **E**, MCF-7 cells (n=2 repeats) were pretreated with a DDD2 (D2) gradient for 24 hours prior to 6TG gradient for an additional 96 hours. **F**, CCRF-CEM cells (n=2) were pre-treated with DDD2 (1 µmol/L) or DDD2 and 0.1/1 µmol/L TH5427 for 24 hours prior to 6TG gradient for an additional 96 hours. **G**, As in **F**, CCRF-CEM cells were pre-treated with DDD2, 5 µmol/L TH7755, or DDD2 and TH7755 for 24 hours prior to adding 6TG. DMSO and DDD2 data from shared experiment in **F** and **G** (n_DDD2_=3, all others n=2). **H**, MCF-7 cells were pretreated with DMSO, TH5427 (5 µmol/L), or DDD2 (1 µmol/L) for 24 hours before addition of 1 µmol/L 6TG for up to 96 hours and Western blotting (representative of n=2 repeats). PARylated and cleaved PARP1 are indicated, while * demarcates non-specific bands. In all cases, statistical significance determined by extra sum-of-squares F-test for EC_50_ difference from control, and error is shown as ± SEM.

We then sought to clarify how NUDT5 affects the thiopurine mechanism-of-action. Thiopurines kill replicating cells by incorporating into DNA, thereby triggering futile mismatch repair (MMR) and toxicity after multiple replication cycles (13,20). NUDT15, another NUDIX enzyme, hydrolyzes active thiopurine triphosphates crucial for clinical efficacy (18–20), so missense polymorphisms and potent NUDT15 inhibitors (NUDT15i) induce thiopurine hypersensitivity (17,30,31). As previously (31), NUDT15i TH7755 sensitized cells to 6TG treatment; however, when combined with DDD2 or NUDT5 shRNA, toxicity was intermediate to either adjuvant alone, suggesting that, while antagonistic, NUDT5 influence is independent of NUDT15 in mediating 6TG activity (**Fig. 2G**, **Supplementary Fig. S3F and G**). Given the 6TG mechanism-of-action, we would also expect less DNA damage-mediated cell death in DDD2-treated cells. To confirm this, we treated cells with NUDT5 compounds and compared the development of 6TG-dependent DNA damage and cell death by following PARP1 modifications (**Fig. 2H**). PARP1 initiates DNA repair through auto-PARylation and is cleaved by pro-apoptotic caspases when damage is too severe (32). As expected, control and TH5427-treated cells demonstrated time-dependent increases in PARylated PARP1 from day 2 and apoptotic cleavage products from day 3. DDD2-treated cells, however, had no changes in PARP1 modifications over this period, aligning with effects seen on overall cell viability. Thus, NUDT5 appears to regulate the toxicity of 6TG metabolites independently of its own catalytic activity or NUDT15, and its depletion decreases the burden of 6TG-mediated DNA damage and cell death.

### NUDT5 mediates the global efficacy of nucleoside analog drugs

After establishing that NUDT5 abundance, but not enzymatic capacity, dictates 6TG efficacy, we then determined if this phenomenon extended to other NA drugs. To this end, we screened a subset of compounds from the MedChemExpress Nucleoside Library by resazurin reduction viability assay (**Fig. 3A**). After triaging cell-impermeable phosphorylated analogs (**Supplementary Table S1**), compounds were added at 0.05, 0.5, or 5 µmol/L to DMSO-, TH5427-, or DDD2-treated cells for 96 hours. To account for incomplete curves and a wide range of toxicity profiles, viability changes were tracked using difference in the area under the curve (ΔAUC) analysis, and analogs that discernably affected control cell viability were prioritized. Of 152 test compounds, 30 evoked viability defects in control cells, and, of those, a striking 24 (80%) were less effective in DDD2 cells, including 6TG and 6MP (**Fig. 3B**, **Supplementary Table S1**). Of the remaining molecules, four were unchanged (N6,N6-dimethyladenosine, floxuridine, 5-azacytidine, adefovir dipivoxil) and two had increased sensitivity when combined with DDD2 (bucladesine and sapacitabine). As before, TH5427 generally had no effect – except for bucladesine and sapacitabine, where cells were sensitized similarly to DDD2.

**Figure 3.**
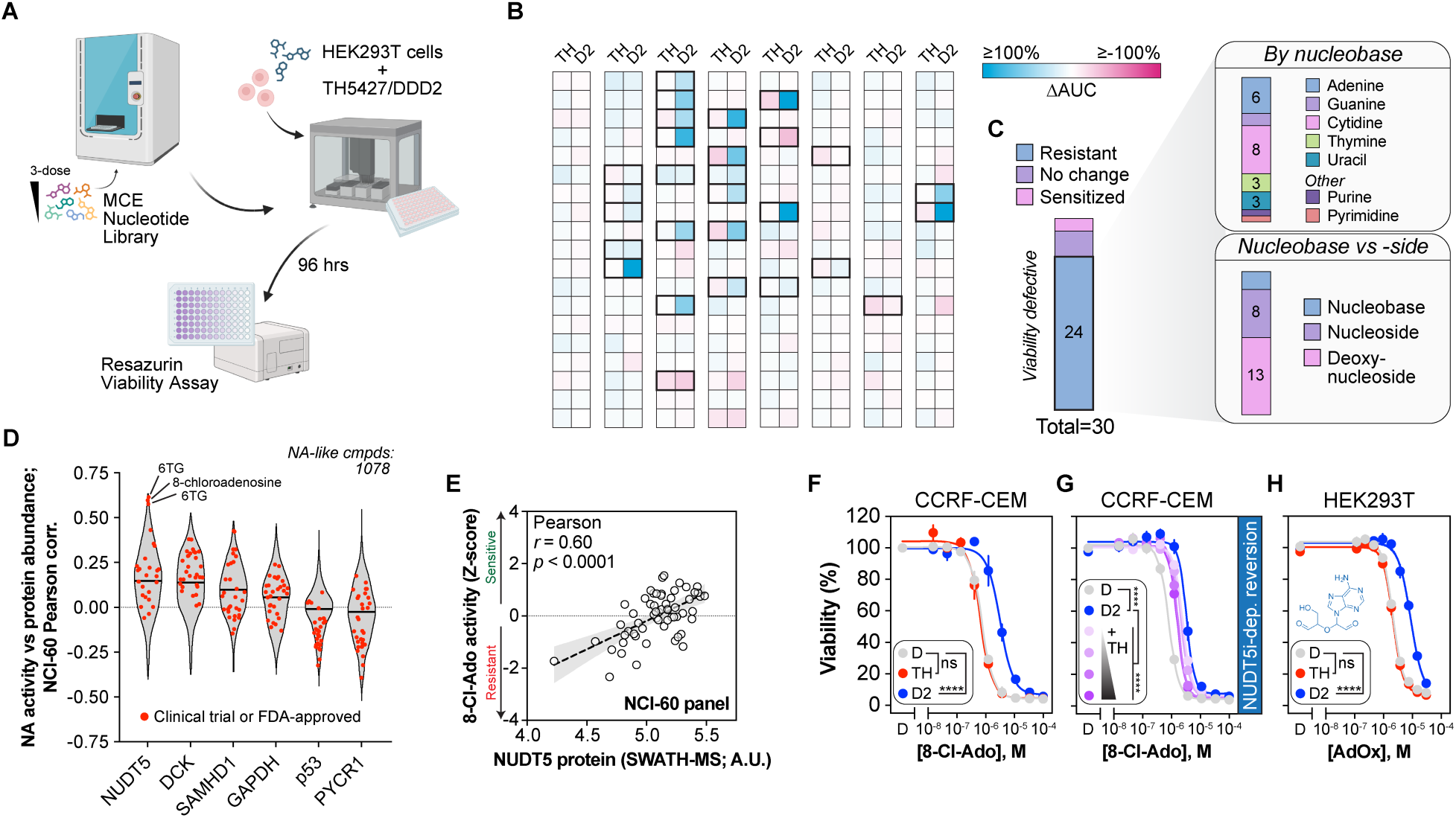
NUDT5 abundance mediates the global efficacy of NA drugs. **A**, Schematic for viability screening of the MCE Nucleotide Library after NUDT5 inhibition or depletion. **B**, NA viability changes following TH5427 or DDD2 depicted as change in area under the curve (ΔAUC). NAs displaying viability defects in control cells are highlighted in bold (n=30). **C**, Breakdown of NA resistance by nucleobase and form. **D**, Activity correlation profiles of 1,078 NA-like compounds in the NCI-60 panel compared among NUDT5, DCK, SAMHD1, GAPDH, p53, and PYCR1. Clinically relevant drugs are highlighted in red. **E**, Pearson correlation of 8-Cl-adenosine (8-Cl-Ado) activity with NUDT5 protein abundance across the NCI-60 panel (*r* = 0.60, *p* <0.0001; linear regression ± 95% CI). **F**, CCRF-CEM cells were pretreated with DMSO, TH5427 (5 µmol/L), or DDD2 (1 µmol/L) for 24 hours and an 8-chloroadenosine (8-Cl-Ado) concentration gradient was added for an additional 96 hours before viability readings (n=2 repeats). **G**, CCRF-CEM cells were treated as in **F** but also in competition with a TH5427 gradient (1 nmol/L to 5 µmol/L; n=2 repeats). **H**, HEK293T cells (n=2 repeats) were pretreated with DMSO, TH5427, or DDD2 before adding an adenosine dialdehyde (AdOx) concentration gradient for an additional 96 hours and viability readings. In all cases, statistical significance determined by two-tailed t test (**E**) or extra sum-of-squares F-test for EC_50_ difference from control (**F**-**H**), and error is ± SEM.

We then categorized the hits to determine trends in DDD2-mediated resistance (**Fig. 3C**). Somewhat surprisingly, there was no obvious tendency towards any one base (or even more broadly towards purines or pyrimidines) or discrimination between nucleobase and nucleoside, although, proportionally, most analogs within the library – and yielding toxicity – were nucleoside forms. For example, both nucleobase 5-fluorouracil (5-FU) and nucleoside 5-fluorouridine toxicity were rescued by DDD2 with 5-fluorouridine being more pronounced (**Supplementary Table S1**). Thus, DDD2-mediated resistance applies generally to cytostatic/cytotoxic NA drugs.

For more comprehensive insights, we again turned to the CellMinerCDB database, which has curated proteomic abundance and drug activity responses for ∼25,000 compounds in the NCI-60 cell lines (28). Triaging of nucleoside-like compounds resulted in 1,078 molecules that could broadly be considered NAs (*i.e.*, containing purine or pyrimidine moieties), and these were used for correlating compound antiproliferative activity with NUDT5 protein expression across the NCI-60 cell lines (**Fig. 3D**). To benchmark correlative observations, data for NUDT5 were compared with enzymes heavily involved in nucleotide and NA drug metabolism (DCK, SAMHD1), as well as proteins indirectly linked to nucleotide metabolism (GAPDH, p53, PYCR1). DCK and SAMHD1 levels generally correlated positively with NA efficacy (median *r* = 0.14 and 0.10, respectively), while GAPDH, p53, and PYCR1 did not. Overall, NUDT5 correlations resembled DCK (median *r* = 0.15), including enrichments in clinical NA drugs. Among the top correlating drug sensitivities were thiopurines and another compound in clinical trials, 8-chloroadenosine (8-Cl-Ado; **Supplementary Table S2**) (33). For reference, NUDT9 mRNA expression did not positively correlate with NA efficacy like NUDT5 mRNA (**Supplementary Fig. S4**, **Supplementary Table S3**).

Closer inspection of 8-Cl-Ado indicated it had a strong positive correlation with NUDT5 protein abundance in the NCI-60 panel (*r* = 0.60, *p* < 0.0001), and, like 6TG, only DDD2 pre-treatment yielded resistance in cells, which could also be reverted by TH5427 titration (**Fig. 3F and G**). Similarly, the screen identified that even NAs that are not obvious substrates for nucleoside kinases, such as adenosine dialdehyde (Ado dialdehyde), a methionine cycle adenosylhomocysteinase (AHCY) inhibitor (34), are rescued by PROTAC-mediated NUDT5 depletion (**Fig. 3H**). Collectively, these data suggest that the abundance of NUDT5 protein, but not enzymatic activity, is required for general efficacy of NAs.

### NUDT5 abundance dictates the efficacy of anti-leukemic NAs in ALL cells

Several NA drugs are front-line therapies for treating cancer, particularly hematological malignancies (35). After establishing that NUDT5 protein regulates the toxicity of many NA drugs, we tested if these effects extended to acute lymphoblastic leukemia (ALL)-approved NA drugs in relevant cell models (CCRF-CEM, Jurkat, MOLT-4, HPB-ALL, MOLT-16, PEER). First, we confirmed that DDD2 degraded NUDT5 in these cells, as hematological cancer cell lines in the NCI-60 panel generally had higher NUDT5 protein levels (**Fig. 4A and B**). Thiopurines, predominantly 6MP, are the backbone of ALL maintenance therapy (36), so we first determined if DDD2 desensitized ALL cells to 6TG and 6MP. NUDT5 depletion, not inhibition, yielded resistance to both thiopurines, with slightly better effect in 6MP-treated cells (**Fig. 4C**, **Supplementary Fig. S5**). HPB-ALL cells, which had inherently high resistance to both thiopurines, had no change in sensitivity, which coincided with poorer depletion of NUDT5 by DDD2 (**Fig. 4A**). We then expanded the cohort to include araG, clofarabine, cytarabine (araC), 8-Cl-Ado, as well as 5-FU (**Fig. 4D**, **Supplementary Fig. S6**). Again, PROTAC pre-treatment yielded resistance to all analogs and TH5427 had no effect, suggesting that NUDT5 abundance may be a relevant predictor of NA drug efficacy in ALL cells.

**Figure 4.**
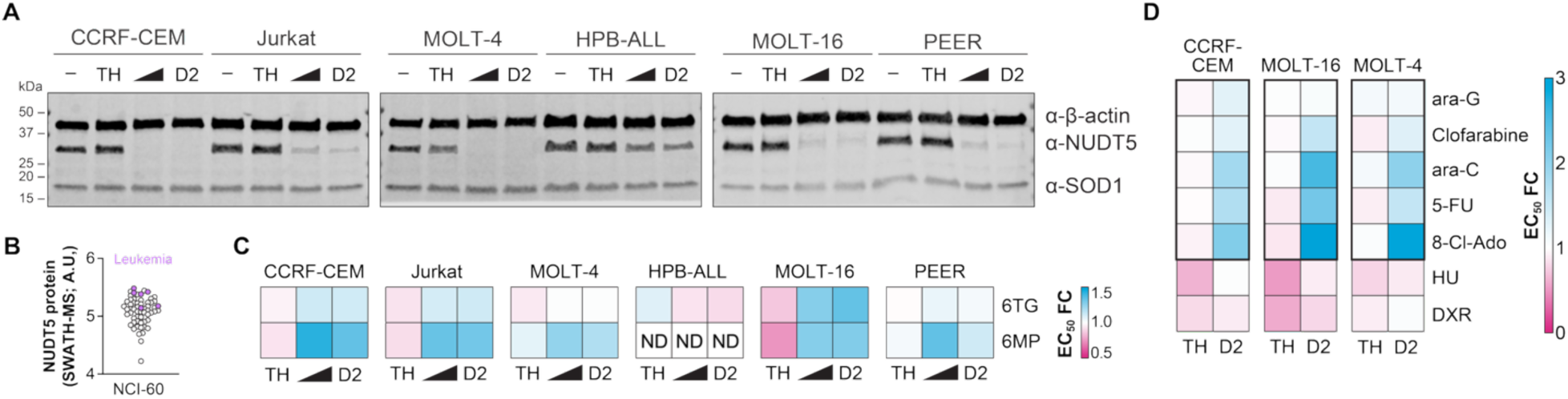
NUDT5 depletion desensitizes ALL cell lines to anti-leukemic NA drugs. **A**, ALL cell lines were incubated with DMSO (–), TH5427 (5 µmol/L), or DDD2 (0.5 or 1 µmol/L) for 24 hours prior to Western blotting to confirm NUDT5 depletion (representative of n=2 experiments). **B**, NUDT5 protein expression levels (SWATH-MS) in the NCI-60 panel with leukemic cell lines highlighted in magenta. **C**, Viability changes expressed as mean EC_50_ fold change (FC) from DMSO control in ALL cell lines pretreated with TH5427 or DDD2 for 24 hours, then a 6TG or 6MP gradient for an additional 96 hours before resazurin measurement (n=3). **D**, CCRF-CEM, MOLT-16, or MOLT-4 cells were treated as in **C** but with the anticancer NAs or chemotherapies indicated (n=3 repeats in **C** and **D**).

Importantly, to this point we had only looked at NUDT5-mediated resistance to NA compounds, so it was possible that observed effects with DDD2 were non-specific. To this end, we included hydroxyurea (HU) and doxorubicin (DXR) titrations alongside our anti-leukemia NA drug panel, as they kill replicating cells *via* alternative mechanisms (**Fig. 4D**, **Supplementary Fig. S6**). HU targets ribonucleotide reductase to induce DNA replication stress *via* dNTP imbalance, while DXR is a DNA intercalator causing double strand breaks by impeding TOP2B progression. Neither compound was affected by DDD2-mediated NUDT5 depletion, whereas TH5427 appeared to slightly sensitize ALL cells to HU. Thus, NUDT5 abundance is not a universal regulator of cell death, rather it intrinsically controls the efficacy of NA drugs.

### NUDT5-depleted cells have inefficient nucleotide salvage

Nucleobase and nucleoside analogs function by piggybacking onto nucleotide salvage metabolic routes, where they are converted to therapeutically actionable forms (12,35). As NAs appeared to be ineffective in NUDT5-depleted cells, it implies that NUDT5 fundamentally affects their activation. Cellular needs for nucleotides are met by the combination of *de novo* and salvage pathways, where *de novo* pathways entail energetically demanding, from-scratch synthesis and salvage pathways center on reclamation of nucleobases and nucleosides (1). The folate cycle supplies one-carbon units to support *de novo* nucleotide synthesis, which is believed to be the main source of nucleotides for proliferating cells, such as cancers (**Fig. 5A**) (2). To determine how NUDT5 depletion impacts viability related to *de novo* nucleotide synthesis, we first starved cells of folate for three days and compared NUDT5 inhibition to degradation (**Fig. 5B**). As expected, folate-deprived cells failed to proliferate (∼15% confluence), while addition of TH5427 had a small, but insignificant, repression of lingering proliferation. However, DDD2-treated cells were consistently more sensitive to folate withdrawal (∼2% confluence). Notably, repletion by various folate intermediates could rescue proliferation in all cases, suggesting folate cycling *per se* is unaffected in NUDT5-depleted cells (**Supplementary Fig. S7A and B**).

**Figure 5.**
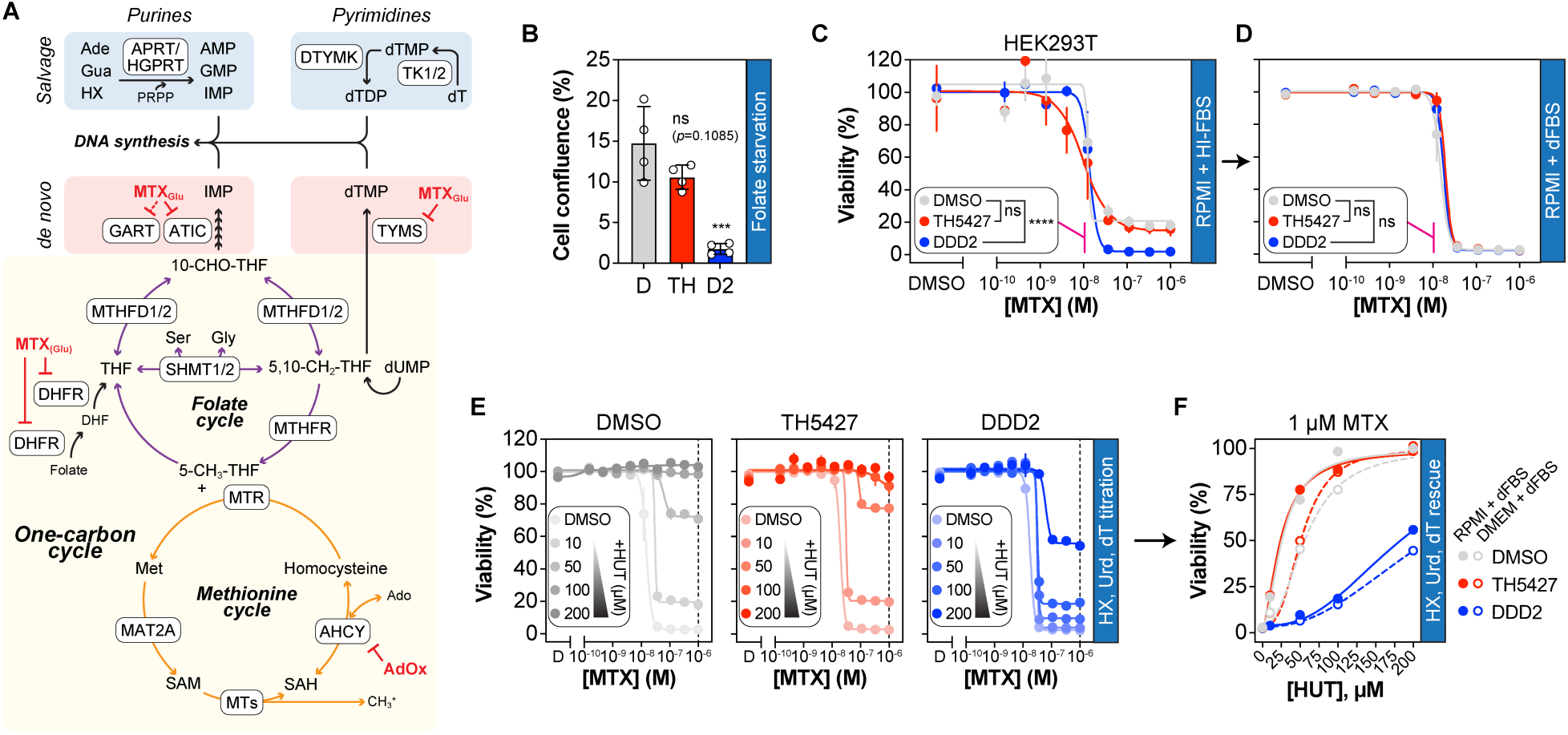
Folate manipulation reveals defective nucleotide salvage in NUDT5 depleted cells. **A**, Schematic depicting the interplay of the one-carbon cycle with *de novo* and salvage nucleotide metabolism needed for DNA synthesis. Relevant inhibitors are indicated in red. **B**, HEK293T cells were pretreated with DMSO (D), TH5427 (TH), or DDD2 (D2) and replated in folate-free medium for 72 hours prior to confluence measurements (CELLCYTE X; n=4; means ± SD). **C**, HEK293T cells were maintained in RPMI + 10% HI-FBS and pretreated with DMSO, TH5427, or DDD2 for 24 hours followed by an MTX gradient for an additional 96 hours (n=2 repeats) before resazurin measurement. **D**, HEK293T cells were maintained in RPMI + 10% dialyzed FBS (dFBS) as in **C** (n=2 repeats). **E**, HEK293T cells were treated as in **D** but with titration of nucleotide precursors (hypoxanthine, uridine, thymidine; HUT) at the indicated concentrations (n=2 repeats). **F**, Cross-sectional representation of mean viability rescue by nucleotide precursors in DMSO-, TH5427-, or DDD2-treated HEK293T cells at 1 µmol/L MTX with comparison between RPMI (filled circles) and DMEM (empty circles) with dFBS (each from n=2 repeats). In all cases, statistical significance determined by one-way ANOVA with multiple comparisons to DMSO (**B**) or extra sum-of-squares F-test for bottom difference compared to DMSO control (**C**, **D**). Means ± SEM are shown in **C**-**E**.

One-carbon metabolism can also be targeted with anti-folates, such as methotrexate (MTX). MTX primarily targets DHFR but, upon polyglutamation, gains nanomolar affinity for AICAR transformylase (ATIC) and thymidylate synthase (TYMS), key cogs in purine and pyrimidine *de novo* synthesis, respectively (**Fig. 5A**) (37). We then subjected TH5427- or DDD2-treated cells to an MTX gradient (**Fig. 5C**, **Supplementary Fig. S7C**). Neither NUDT5 inhibition nor depletion meaningfully shifted the viability curve in response to MTX; however, DMSO- and TH5427-treated cells displayed a viability plateau (∼15-20%), while DDD2 cells were near 0%. This effect notably mirrored folate deprivation (**Fig. 5B**, **Supplementary Fig. S7B**), indicating that NUDT5 enables residual survival in folate deprived cells.

Anti-folates block *de novo* nucleotide production, but cells can circumvent this by using nucleotide salvage to sustain DNA synthesis (38). Typical cell culture fetal bovine serum (FBS) contains nucleosides, which desensitize cells to MTX, but these can be effectively removed by dialysis to revert the effect (38). When we changed to dialyzed FBS, the MTX viability curves for all three pretreatments bottomed out and were indistinguishable from each other (**Fig. 5D**, **Supplementary Fig. S7D**). This suggested that NUDT5 degradation specifically suppressed the contributions of nucleotide salvage to remaining cell survival. To test this in practice, we again subjected all three pretreatments to an MTX gradient in dialyzed FBS and supplemented cells with increasing concentrations of nucleotide precursors (hypoxanthine [HX], uridine [Urd], and thymidine [dT]) to induce nucleotide salvage (**Fig. 5E and F**, **Supplementary Fig. S7C and D**). Control and TH5427-pretreated cells were similarly rescued – fully reverting MTX toxicity at ∼100 µmol/L concentrations – however, DDD2 severely blunted MTX rescue by a factor of up to 10-fold, irrespective of whether RPMI or DMEM media formulations were used. Like previous studies (38), titration of individual precursors was not enough to overcome MTX toxicity in any case (**Supplementary Fig. S7E**). Collectively, these results suggest that low NUDT5 abundance dampens nucleotide salvage capabilities leading to ineffective reversal of MTX toxicity.

### NUDT5 depletion locks cells into de novo purine biosynthesis

To this point, our results suggested that NUDT5 protein *per se*, and not its enzymatic activity, are required for the global efficacy of NA drugs, which may be explained by inefficient nucleotide salvage in NUDT5-depleted cells. We then turned to global metabolomics analyses to corroborate these findings by monitoring NUDT5-dependent metabolic changes over 24 hours, as well as the influence of 6TG on these changes (**Fig. 6A**). Metabolites detected *via* LC-MS analysis were generally unchanged irrespective of NUDT5 targeting approach or 6TG addition with some notable exceptions (**Fig. 6B**, **Supplementary Fig. S8A**, **Supplementary Table S4**). Somewhat surprisingly, we saw that 5-aminoimidazole-4-carboxamide ribonucleotide (AICAR), a key intermediate of *de novo* purine biosynthesis (DNPB) (1), was suppressed following addition of 6TG, suggesting that DNPB was restricted (**Fig. 6C**). While 6MP intermediates are known to block DNPB *via* amidophosphoribosyltransferase (PPAT) inhibition as part of its antiproliferative effects (39) – 6TG metabolites are also capable of repressing DNPB, although its anticancer effects are primarily attributed to nucleic acid incorporation (40). TH5427 addition resulted in a decreased basal AICAR abundance that was exacerbated upon 6TG addition, while, strikingly, DDD2 slightly increased basal AICAR levels and was completely unaffected by 6TG. Both TH5427 and DDD2 also increased the abundance of amino acids (AAs), with DDD2 having faster dynamics and a stronger enrichment for aromatic amino acids (phenylalanine, tyrosine, tryptophan) and was not influenced by 6TG (**Fig. 6D**). We also observed this for amino acids detected by GC-MS, albeit with worse resolution (**Fig. 6E**, **Supplementary Fig. S8B**, **Supplementary Table S4**). This phenomenon has been observed previously by Hoxhaj *et al.* when treating cells with DNPB inhibitors and was attributed to collateral effects of mTORC1 inhibition by purine starvation (41). Ribose was also selectively elevated in DDD2-treated cells by GC-MS but not in control or TH5427-treated counterparts (**Fig. 6F**). Free ribose can be salvaged from serum or generated by myriad ribohydrolases *via* reversible conversion of nucleosides to bases and directly converted to R5P by ribokinase (RBKS) for generating PRPP and ribose-1-phosphate, thereby bypassing the more costly oxidative PPP (42,43). Thus, NUDT5 depletion appears to lock cells into DNPB, even when disadvantageous, and suppresses utilization of ribose, a key salvage metabolite.

**Figure 6.**
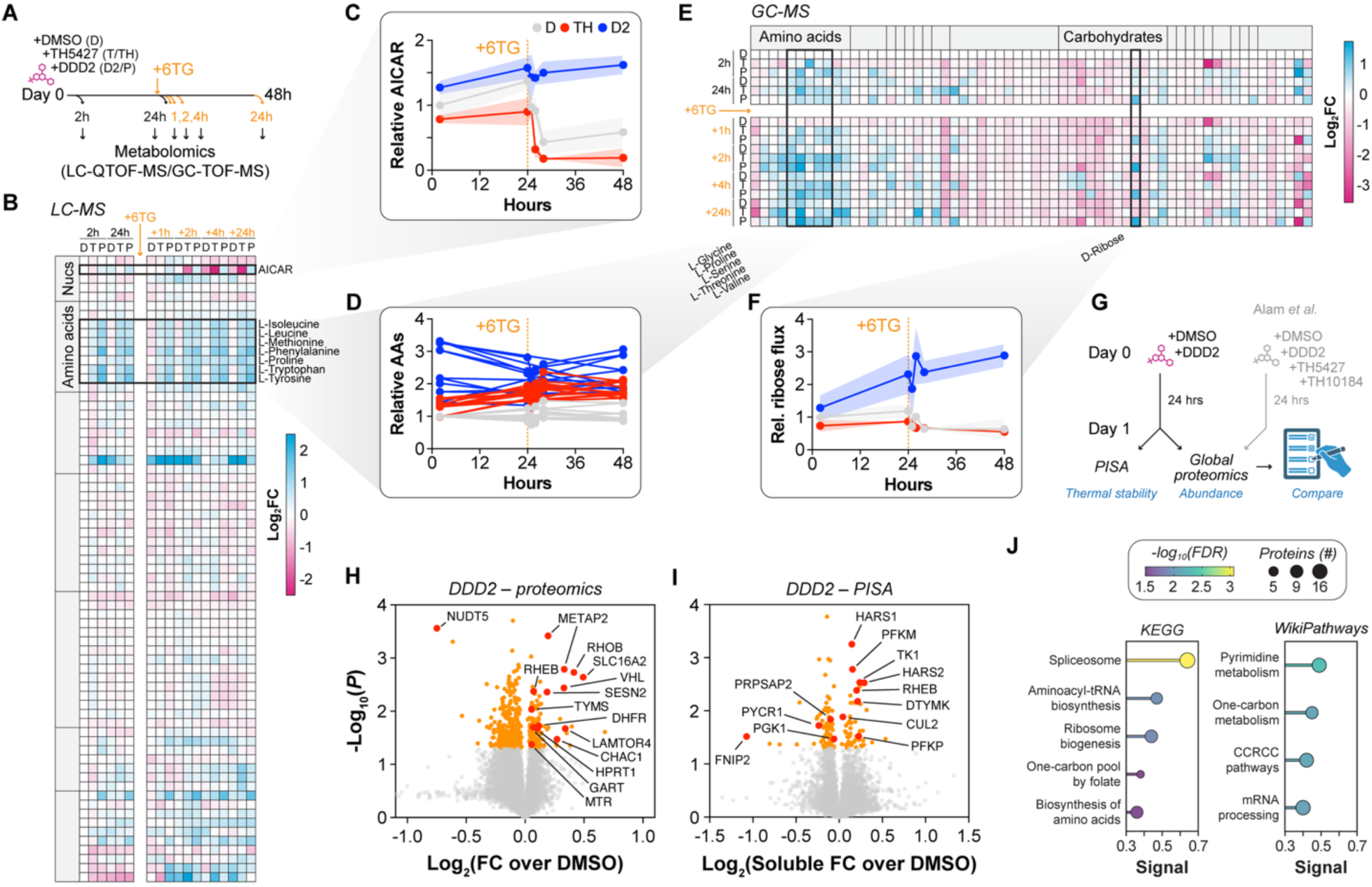
Multi-omics analysis suggests NUDT5-depleted cells are locked into *de novo* purine synthesis. **A**, Schematic of longitudinal metabolomics analysis of DMSO (D), TH5427-(T/TH), or DDD2 (D2/P)-pre-treated cells ± 1 µmol/L 6TG (orange – time since addition) over 48 hours. **B**, Heatmap of log_2_FC metabolite abundance changes by LC-QTOF-MS. All changes relative to DMSO, 2 hours and are means from n=3 replicates. Notable metabolites are highlighted. **C**, Relative abundance of AICAR and individual amino acids (AAs; **D**) in DMSO, TH5427, or DDD2-treated cells over time with 6TG addition indicated. Means shown ± SD. **E**, Heatmap of log_2_FC metabolite abundance changes by GC-TOF-MS. All changes relative to DMSO, 2 hours and are means from n=3 replicates. Notable metabolites are highlighted. **F**, Relative abundance of D-ribose in DMSO, TH5427, or DDD2-treated cells over time with 6TG addition indicated. Means shown ± SD. **G**, Proteomics strategy overview. Cells were incubated with DMSO or DDD2 for 24 hours prior to global proteomics or PISA analysis. Results are compared with global proteomics results from Alam *et al.* with TH5427 and DDD2 (ProteomeXchange ID: PXD065527, (29)). Part of this figure was made in BioRender. **H**, Global proteomics volcano plot of protein abundance changes after 24 hours DDD2. Significant hits shown in orange and highlighted hits shown in red. Means from n=4 experiments shown. **I**, PISA volcano plots from cells treated with DDD2 for 24 hours and shown as in **H**. **J**, Lollipop plot representations of pathway enrichment analyses for combined global proteomics and PISA hits for DDD2 benchmarked to KEGG and WikiPathways databases. Enrichments shown as signal, while -log_10_(FDR) and number of proteins are shown by color gradient and circle size, respectively.

Notably, despite significant fluctuations in AICAR and ribose, the abundance of purine (AMP, IMP) and pyrimidine nucleotides (UMP) remained stable – although at two hours TH5427-treated cells had decreased AMP, perhaps reflecting the acute inhibition of ADP-ribose turnover by NUDT5 (**Fig. 6B and E**). This suggested that overall nucleotide precursor pools were not significantly affected, in line with the observation that neither NUDT5 inhibition nor depletion adversely affect proliferation.

### Proteome-wide assessment of NUDT5-targeting small molecules

Alongside metabolic profiling, we also performed proteomics analyses to better understand how metabolic changes were reflected at the protein level to compare NUDT5 inhibition and depletion after 24 hours. To facilitate analysis, an earlier dataset comparing the NUDT5 inhibitors TH5427 and TH10184 to DDD2 was included alongside a second DDD2 proteomic analysis reported here (**Supplementary Fig. S9A-C**). In the second analysis, DDD2-treated cells were also assessed for global thermal stability changes using the proteome integral solubility alteration (PISA) assay, which can detect biophysical changes caused by *e.g.* ligand binding, post-translational modifications, and changes in protein complex dynamics (44) (**Fig. 6G**). In total, 9751 proteins were quantified (≥ 2 peptides), while the relative abundance of 446 and 131 proteins were significantly changed generally and by PISA after DDD2 treatment, respectively (**Supplementary Table S5 and S6**). As previously (29), DDD2 significantly degraded NUDT5 and appeared to slightly increase the abundance of VHL and stabilize CUL2, the co-opted E3 ligase complex mediating DDD2-dependent proteolysis (**Fig. 6H and I**). Some of the most significantly changed proteins included SLC family members (SLC47A1, SLC16A2, SLC7A2), mTORC1-related amino acid sensing proteins (SESN2, LAMTOR4, FNIP2, RHEB), ATF4-dependent genes (CHAC1, tRNA synthetases), one-carbon and pyrimidine metabolic enzymes (DHFR, TYMS, GART, TK1, DTYMK), glycolytic enzymes (PFKM, PFKP, PGK1) and the N-terminal methionine aminopeptidase, METAP2. TH5427 had significant overlap with NUDT5 inhibitor, TH10184, (∼40%), and 120 proteins overlapped with DDD2 enrichments within the first experiment. However, there was little similarity when including comparisons to the second set of DDD2 proteomics (39 DDD2-common and 11 overall common proteins; **Supplementary Fig. S9A-C**).

We then performed a functional enrichment analysis of the combined results for DDD2 using STRING DB to understand which systems are altered in DDD2-treated cells (**Fig. 6J**). Benchmarking to KEGG and WikiPathways indicated some enrichment in the one-carbon and pyrimidine metabolism pathways, among others related to mRNA processing/spliceosome and ribosome biogenesis, in line with our observations of perturbed nucleotide metabolism in DDD2-treated cells. Nonetheless, like the metabolomics data, NUDT5 depletion was not decidedly different from inhibition, where many proteins, such as TYMS and MTR, were regulated in the same manner in both treatments (**Supplementary Fig. S9C**).

### NUDT5 interacts with PPAT, the rate-limiting enzyme of DNPB

Our multi-omics approach to deconvolute the mechanisms by which NUDT5 degradation, not inhibition, causes NA drug resistance were largely inconclusive – save for the persistent flux through DNPB seen in NUDT5-depleted cells. Thus, we investigated potential protein-protein interactions (PPIs) that could explain observed effects. Intriguingly, curated data from large-scale interactome experiments has indicated that PPAT, the first enzyme in DNPB, is a potential NUDT5 interactor, as well as pyrroline-5-carboxylate reductase 1 and 2 (PYCR1, PYCR2) (45). We sought to confirm these observations by expressing N- or C-terminal GFP fusions of NUDT5 and co-immunoprecipitating complexes by GFP-Trap (**Fig. 7A**). This approach was chosen to ensure a clean pull-down of the NUDT5 bait and gain insights to complex orientation. As expected, the GFP-tagged NUDT5 co-immunoprecipitated with endogenous NUDT5, which forms a homodimer (26). Both PYCR1 and PYCR2 were also abundantly complexed with NUDT5 irrespective of tag orientation, while PPAT interacted with NUDT5 only when tagged on the N-terminus. None were present with GFP pull-down alone, but all three also co-immunoprecipitated with endogenous NUDT5, thereby confirming their interaction (**Supplementary Fig. S10A**).

**Figure 7.**
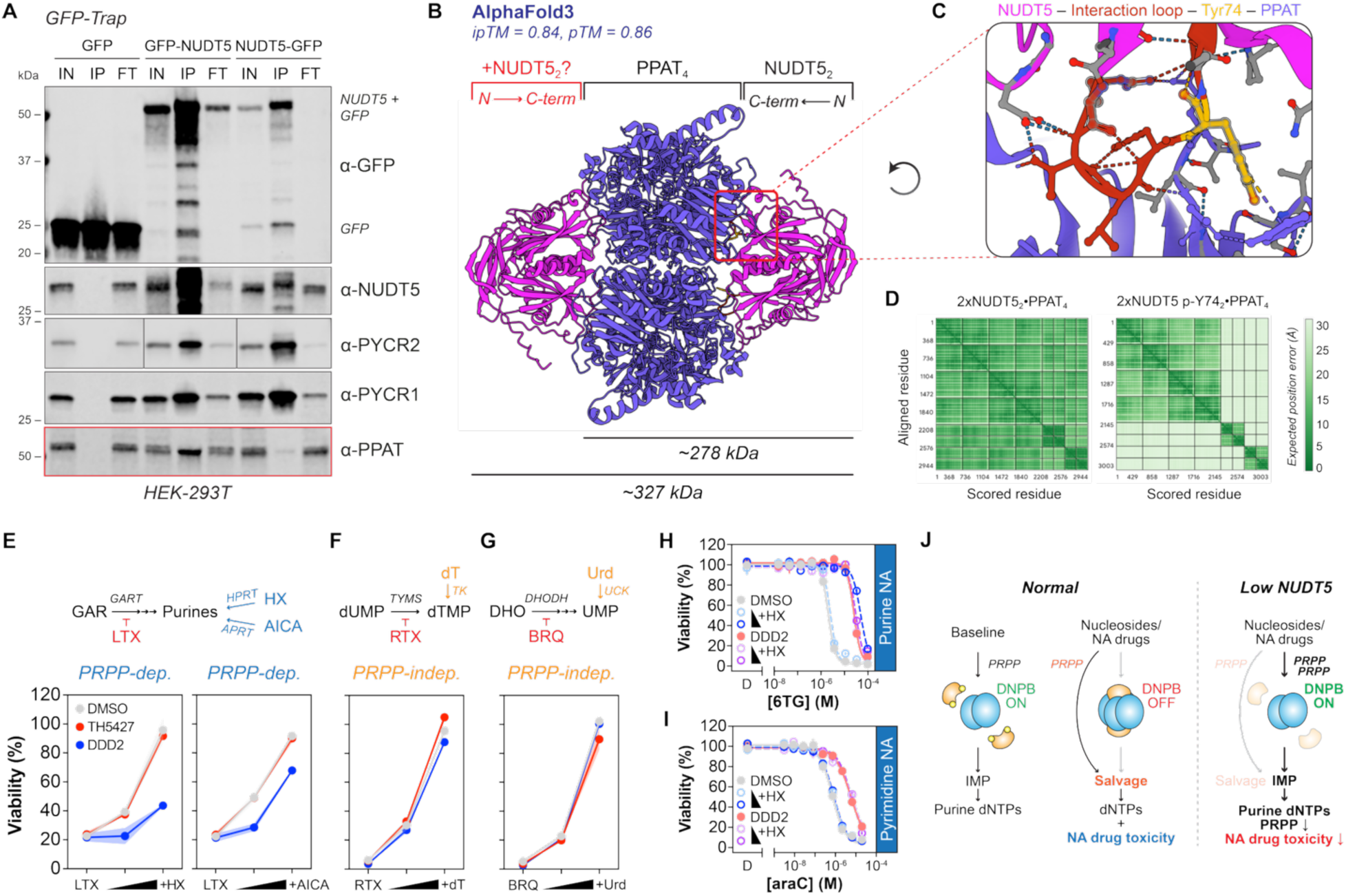
NUDT5 interacts with PPAT to regulate DNPB and NA drug efficacy through PRPP misallocation. **A**, GFP-Trap co-immunoprecipitation analysis of GFP-tagged NUDT5 on the N-or C-terminus. Representative experiment shown from n=2. **B**, AlphaFold 3 complex modeling between NUDT5 (magenta) and PPAT (purple). Prediction confidence scores are provided for both (NUDT5_2_)_2_•PPAT_4_ and (NUDT5_2_)_4_•PPAT_4_ structures, as well as estimated molecular weights (kDa). **C**, Zoomed inset of a proposed PPAT interaction loop spanning residues 70-76 (red) on NUDT5, with potential phosphorylation site, Tyr74, highlighted in yellow. **D**, Per-residue PAE confidence scores for predicted AlphaFold structures for NUDT5 WT (2xNUDT5_2_•PPAT_4_) and phospho-Tyr74 (2x NUDT5 p-Y74_2_•PPAT_4_). **E**, HEK293T cells were grown in DMEM with dFBS before pre-treatment with DMSO (1% v/v final; grey), 5 µmol/L TH5427 (red), or 1 µmol/L DDD2 (blue) for 24 hours, followed by 111 nmol/L lometrexol (LTX) ± 10/100 µmol/L hypoxanthine (HX) or ± 100/150 µmol/L 5-amino-3H-imidazole-4-carboxamide (AICA) for an additional 96 hours. Viability was then determined by resazurin reduction assay. **F**, Cells were treated as in **E** but with 1.11 µmol/L raltitrexed (RTX) ± 2/10 µmol/L thymidine (dT). **G**, Cells were treated as in **E** but with 1.11 µmol/L BRQ ± 10/100 µmol/L uridine (Urd). Data shown in **E**-**G** as means ± SEM from n=2-3 independent experiments. **H**, HEK293T cells were grown in DMEM with dFBS before pre-treatment with DMSO (grey) or DDD2 (blue) for 24 hours, followed by 6TG or araC (**I**) gradient ± 10/100 µmol/L HX (blue and purple, respectively) for an additional 96 prior to viability assessment by resazurin assay. **J**, Proposed model of PPAT (blue) regulation by NUDT5 (orange) depending on phosphorylation of Tyr74 (yellow) to control DNPB flux. NA drug efficacy is driven by PRPP liberated from NUDT5-mediated inactivation of DNPB via PPAT. When NUDT5 is depleted, DNPB is persistent, consuming PRPP and thereby inactivating NA drugs.

As PPAT is the first and committal step of DNPB, we took a closer look at its interaction with NUDT5 to potentially explain the observed DNPB persistence and NA drug resistance seen in DDD2-treated cells. PPAT is itself a homomultimeric protein known to exist in at least two states: an active dimeric state (PPAT_2_) and an inactive tetrameric state (PPAT_4_) (46). These activity states are sensitive to metabolic feedback, where PRPP addition shifts the complex to the active conformation (high DNPB activity), while the addition of purine nucleosides (high salvage activity) transitions to the larger, inactive form (46). Nonetheless, structures of human PPAT have not been resolved, presumably due to its labile FeS clusters (47), thus we turned to the AlphaFold Server to model both potential PPAT and NUDT5-PPAT complexes (48). As expected, PPAT was modeled with high accuracy in both the dimeric (active) and tetrameric (inactive) states (**Supplementary Fig. S10B and C**), where the PPAT_4_ barrel structure was like homologous purF structures reported for *B. subtilis* (49) and *E. coli* (50) (**Supplementary Fig. S10D**).

We then turned to NUDT5-PPAT complexes, which could also inform on whether this was a direct interaction or *via* possible intermediaries. Low confidence predictions were observed with one (ipTM = 0.57) or two (ipTM = 0.68) NUDT5 dimers (NUDT5_2_) modelled with the active PPAT_2_ conformation (**Supplementary Fig. S10E**). Conversely, we obtained high accuracy and high confidence models of one or two NUDT5_2_ capping the ends of the PPAT_4_ barrel (pTM and ipTM for both 0.84 and 0.86, respectively, overall high pIDDT, PAE; **Fig. 7B**). Notably, this model aligns with our observations that C-terminal GFP tagging disrupts PPAT binding to NUDT5, presumably due to steric displacement of the PPAT_4_ barrel. A small loop within the NUDIX fold beta sheet, comprising residues 70 to 76, appeared to be critical to interaction with PPAT residues Gln26, Asp28, Tyr83, and Arg363 in the (NUDT5_2_)_n_•PPAT_4_ models (**Fig. 7C**). Tyrosine 74 (Tyr74) is annotated as a potential phosphorylation site (51), which may be capable of disrupting the NUDT5-PPAT complex. Indeed, when we modelled the complex with phospho-Tyr74, the accuracy and PAE confidence metrics suggested that it was effectively disrupted when compared with wild-type NUDT5 (pTM = 0.59 and ipTM = 0.53, respectively; **Fig. 7D**, **Supplementary Fig. S10F**). These data collectively suggest that NUDT5 directly interacts with PPAT in its inactive (PPAT_4_) state *via* an interaction loop (res 70-76) conditional upon Tyr74 phosphorylation.

### NUDT5 depletion selectively impairs PRPP-mediated nucleotide biosynthesis

PPAT initiates DNPB by building onto PRPP, and, like UMPS in the *de novo* pyrimidine biosynthesis pathway, is a primary consumer of this activated ribose scaffold (1). Numerous studies have suggested that the PRPP pool is a limited commodity that is consumed by competing enzymes (5–8,11). If NUDT5 protein is regulating PPAT activity, it would be conceivable that PRPP-dependent processes are impaired upon NUDT5 depletion, *ie* when PPAT is constitutively active. To test this, we revisited antimetabolite therapeutics that can be effectively rescued by defined nucleotide precursors: lometrexol (LTX), which inhibits DNPB *via* GART; raltitrexed (RTX), which inhibits *de novo* dTMP synthesis *via* TYMS; and brequinar (BRQ), which inhibits *de novo* pyrimidine biosynthesis *via* DHODH (**Fig. 7E-G**, **Supplementary Fig. 11**). As before, cells were cultured in dFBS to minimize the influence of serum-derived nucleotide salvage. In all cases, control or TH5427-treated cells were effectively rescued by all nucleotide precursors, whereas DDD2 cells had impaired rescue by PRPP-dependent metabolites (hypoxanthine [HPRT], AICA [APRT] rescue of LTX) but not PRPP-independent processes (thymidine [TK] rescue of RTX; uridine [UCK] rescue of BRQ). Notably, this effect was irrespective of whether rescue agents bypassed *via* the salvage pathway or downstream of blockades in *de novo* pathways – such as AICA conversion to AICAR by APRT just downstream of PPAT and GART, the target of LTX.

DNPB overactivity can also conceivably impair purine NA drug efficacy by increasing endogenous purines that compete for salvage enzymes (52). When we tested purine NA drug toxicity in regular and dialyzed FBS to understand the contribution of salvageable purines to toxicity, we saw slight sensitization when using dialyzed FBS in both control and NUDT5-depleted cells (**Supplementary Fig. 12**). Likewise, hypoxanthine supplementation desensitized control cells to 6TG, but did not do so in DDD2-treated cells, nor did it contribute to NUDT5-dependent cytarabine resistance, arguing that increased availability of salvageable purine precursors is not driving the effect (**Fig. 7H and I**, **Supplementary Fig 12**). Collectively, these findings suggest that NUDT5 depletion specifically impairs PRPP-dependent processes, in line with constitutively active PPAT exhausting PRPP pools to dysregulate nucleotide metabolism homeostasis.

### PPAT protein abundance inversely correlates with global NA drug efficacy

DNPB expression and activity level can dictate responses to NA drugs by modulating both purine and PRPP pools (5,8,10,11,52,53). We again revisited the CellMinerCDB datasets to profile the efficacy of 1,078 NA-like compounds in relation to the abundance of DNPB enzyme mRNA or protein expression (**Supplementary Fig. S13A and B**, **Supplementary Table S2 and S3**). Further, we could assess their metabolic similarity to NUDT5 by plotting per-compound Pearson correlation coefficients, where high correlation would be deemed nearly identical to NUDT5 (**Supplementary Fig. S13C**). DNPB proteins had a strong positive correlation with NUDT5-NA compound associations (mean *r* = 0.64), which extended to most enzymes involved in *de novo* pyrimidine biosynthesis and nucleotide salvage but not non-nucleotide metabolism proteins, such as p53 and PYCR1, nor paralogous NUDT9 (**Supplementary Fig. S13D-H**). The only enzyme that upended this trend was PPAT, where thiopurine sensitivity correlations were inverted (**Supplementary Fig. S13A, D, and E**). Notably, however, all mRNA associations, including PPAT, were positively correlated to NUDT5, suggesting that the PPAT protein divergence is post-translational (**Supplementary Fig. S13B, D, and E**). Thus, the overall efficacy of salvage-promoting NA drugs is worse with higher PPAT protein levels, consistent with the notion that it is a gatekeeper of DNPB and NA drug efficacy that is potentially regulated by NUDT5.

## Discussion

Exploiting cancer cells’ demand for DNA synthesis with anti-metabolites has been one of the most effective therapeutic strategies since their emergence in the 1940s (2). The reliance on enhanced glycolytic flux *via* the Warburg effect has enabled successful use of anti-folates to target *de novo* nucleotide synthesis. While blockade of nucleotide salvage has not historically been an anticancer strategy, NA drugs utilize these pathways to form active metabolites and target active DNA synthesis in proliferating cancer cells (2,35). Still, their complexity means that new factors regulating their efficacy continually emerge, most of which directly metabolize these drugs to impart their effects.

Here, we report that NUDT5 indirectly affects the anticancer efficacy of thiopurines *via* regulating DNPB activity. Strikingly, only depletion of NUDT5 protein, but not catalytic inhibition, evoked persistent DNPB activity that rendered thiopurines up to 10-fold less effective. This effect was titratable based on NUDT5 abundance and appeared to directly correlate with cell line expression profiles. NUDT5 interacts with the rate-limiting DNPB enzyme, PPAT, collectively suggesting that it suppresses PPAT activity to regulate flux through DNPB. Considering the emerging roles of nucleotide salvage in cancer progression and therapeutic resistance (2,54), targeting NUDT5 *in vivo* to modulate this balance could be therapeutically advantageous (55).

Our results complement other recent studies that converged on this mechanism through different approaches (53,56–59). Further, we significantly expand NUDT5 influence to global NA drug efficacy, as NUDT5 depletion rendered cells resistant to most cytotoxic NA drugs tested, including pyrimidine and nucleoside analogs, suggesting that a common resource is involved. PRPP is a strong candidate, as it is a limited metabolite competitively utilized by all human nucleotide synthesis pathways except pyrimidine salvage, which relies on conversion to nucleotides by UPP1/2 and TK1/2 (2,9,11). To this end, we saw impaired PRPP-dependent rescue of antimetabolite therapies, but PRPP-independent rescue was unaffected. This possibility is further supported by pooling of D-ribose, a salvageable PRPP precursor, in NUDT5-depleted cells – presumably by increased ribohydrolase activity, which could be used to salvage PRPP *via* ribokinase under metabolic stress. DNPB is a major PRPP consumer and its blockade or overactivation significantly increases PRPP pools (8,10,11,52). Critically, liberated PRPP can be used for synergistic activation of thiopurines and 5-FU *via* HPRT and UMPS, respectively (5–8,10). Alternatively, DNPB overactivation by feedback inhibition-defective PRPS1 mutants can drive thiopurine relapse by over-producing purines that out-compete prodrug activation by HPRT (10,52). Our findings that 6TG toxicity is rescued by hypoxanthine supplementation in control cells but not DDD2-treated counterparts would suggest that PRPP misallocation is a predominant factor in NA drug resistance due to high DNPB activity. This is further supported by our anti-folate data suggesting that nucleotide precursors are poorly utilized, even though purines should be abundant in NUDT5-depleted cells.

Similar resistance could conceivably extend to pyrimidine analogs by affecting PRPP availability for *de novo* pyrimidine biosynthesis and contribute to imbalanced nucleotide pools (7,8,11). This nucleotide pool imbalance was likely masked in our metabolomics analysis by salvage of precursors from non-dialyzed FBS but is likely overall insignificant, as conditional NUDT5 depletion did not incur gross viability defects in our hands. Presumably, UMPS-mediated synthesis of UMP from orotate is also impaired in NUDT5-depleted cells, as was also suggested in another study (56), but we were unable to rescue brequinar-dependent toxicity with orotate supplementation, which may relate to its poor cellular uptake (60). Nonetheless, we saw upregulation of TYMS and stabilization of TK1 and DTYMK in NUDT5-depleted cells, consistent with thymidine salvage being activated. Likewise, mimicry of DNPB activation by hypoxanthine supplementation had no effect on cytarabine toxicity in either control or NUDT5-depleted cells, arguing that pyrimidine analog desensitization is not driven by increased purine availability alone. The nature and regulation of the NUDT5-PPAT complex will warrant further investigation.

It is known that PRPP promotes DNPB by prompting formation of active PPAT dimers, whereas purine nucleotides induce the feedback-mediated inactive tetramer (46). Our AlphaFold modeling of potential NUDT5-PPAT structures suggests that NUDT5 preferentially complexes with PPAT in its inactive, tetrameric form, while a putative interaction loop within NUDT5 implicates phosphorylation of Tyr74 regulates complex formation. This is consistent with NUDT5 depletion apparently locking cells into DNPB synthesis and NUDT5 abundance proportionally affecting it. Indeed, related studies have shown that this complex is dependent on a functional interaction loop while being promoted by adenosine and disrupted by PRPP (53,56,59), which suggest the enticing possibility that NUDT5 directly controls DNPB activity *via* PPAT oligomerization.

Curiously, the contribution of NUDT5 inhibition appears to be minimal despite its ADPr hydrolysis products, AMP and R5P, being closely associated with PPAT feedback regulation – although, we note that NUDT5 inhibition additively repressed DNPB with 6TG. These divergent outcomes would suggest that NUDT5 is a sensor that finetunes DNPB based on metabolic cues – such as the availability of salvageable nucleotide precursors and NA drugs, which activate salvage synthesis (61) – a possibility also raised by others that will require further study (53,59). Likewise, the contribution of PYCR1/2 interactions to observed effects also warrant further investigation given their importance to cancer metabolism (55,62,63). Nonetheless, our findings strongly position NUDT5 as a critical determinant of nucleotide metabolic balance and NA drug efficacy with clinical ramifications.

## Methods

### Cell lines and culturing conditions

HEK293T (293T; CRL-3216), MCF-7 (HTB-22), Jurkat (TIB-202), CCRF-CEM (CCL-119), HL-60 (CCL-240) and MOLT-4 (CRL-1582) cell lines were obtained from the American Type Culture Collection (ATCC, Manassass, VA, USA). MOLT-16 (ACC 29), HPB-ALL (ACC 483), and PEER (ACC 6) cell lines were obtained from the German Collection of Microorganisms and Cell Cultures (DSMZ, Germany). HEK293T and MCF-7 cells were cultured in DMEM high glucose, GlutaMAX medium (Thermo Fisher Scientific), and Jurkat, CCRF-CEM, Molt4, HPB-ALL, Molt16, and PEER cells were cultured in RPMI, GlutaMAX medium (Thermo Fisher Scientific), unless otherwise stated. All media were supplemented with 10% heat-inactivated fetal bovine serum (FBS; Thermo Fisher Scientific) and penicillin/streptomycin, unless otherwise stated. Cell cultures were maintained at 37 °C with 5% CO_2_ in a humidified incubator. Purchased cell lines were originally authenticated by the ATCC and DSMZ (STR profiling), while MOLT-4, MOLT-16, HPB-ALL, PEER, Jurkat, and CCRF-CEM lines were re-validated by STR profiling (Microsynth AG) – no further authentication was performed for other cell lines. The cells were routinely screened for mycoplasma using the MycoAlert kit (Lonza Bioscience) and, except for Jurkat, CCRF-CEM, HL-60, HPB-ALL, and PEER, none were listed as misidentified on ICLAC or known to be cross-contaminated.

### Antibodies and chemicals

Anti-GFP (rabbit, cat. #sc-8334), anti-NUDT5 (mouse, clone E-4, cat. # sc-398644), anti-PARP1 (mouse, clone F-2, cat. #sc-8007), and anti-SOD1 (mouse, clone G-11, cat. #sc-17767) were obtained from Santa Cruz Biotechnology. anti-NUDT5 (rabbit polyclonal) was generated in-house as previously described (26). anti-β-actin (mouse, clone AC-15, cat. #ab6276) and anti-NUDT5 (rabbit, clone EPR7734, cat. # ab129172) were purchased from Abcam. Anti-PARP1 (rabbit, clone 3N19, cat. #80174-1-RR) was purchased from ProteinTech. Anti-PYCR1 (rabbit, cat. # A304-836A) was purchased from Bethyl Laboratories. Anti-PYCR2 (mouse, clone 4A10, cat # GTX83752) was purchased from GeneTex. Anti-PPAT (rabbit, cat # PA5-27770) was purchased from Thermo Fisher Scientific. Donkey anti-mouse IgG IRDye 680RD (cat. #925-68072, lot #D20803-13) and goat anti-rabbit IgG IRDye 800CW (cat. #925-32211, lot #D21109-25) were purchased from LICORbio.

Doxycycline hydrochloride (Sigma-Aldrich) was dissolved in MilliQ water (2 mg/mL) and used at indicated concentrations. Resazurin sodium salt (Sigma Aldrich) was dissolved in PBS to 1 mg/mL and filtered prior to aliquoting. AraG, fluorouracil (5-FU), clofarabine, cytarabine (araC), hydroxyurea (HU) and doxorubicin (DXR) were purchased from Sigma Aldrich. 6-thioguanine (6TG), 8-chloro-adenosine (8-Cl-Ado), adenosine dialdehyde (AdOx), methotrexate, raltitrexed, hypoxanthine (HX), uridine (Urd), thymidine (dT), folic acid, 5-formyl-THF, and 5-methyl-THF were purchased from MedChemExpress and dissolved in DMSO. 6-mercaptopurine (6-MP) and 6TG were also purchased from Enamine and dissolved in DMSO. TH5427 hydrochloride was purchased from Tocris and dissolved in DMSO (10 mmol/L). Synthesis of DDD2 was described previously (29) and dissolved in DMSO (10 mmol/L). The nucleoside analog library (HY-L044) used in the viability screen was from MedChemExpress (more details found in the screening methodology).

### Recombinant DNA cloning

#### Cloning procedures

shRNA expression plasmids were generated by primer annealing and standard BshTI/EcoRI restriction digestion/ligation into pRSITEP-U6Tet-(sh)-EF1-TetRep-P2A-Puro-P2A-RFP670, as before (64), using target sequences from The RNAi Consortium (TRC). The following primers were used: shNT – 5’-CCGGCCTAAGGTTAAGTCGCCCTCGCTCGAGCGAGGGCGACT-TAACCTTAGGTTTTTG -3’ (F), 5’-AATTCAAAAACCTAAGGTTAAGTCGCCCTCGCTC-GAGCGAGGGCGACTTAACCTTAGG -3’ (R) (64); shNUDT5-1 – 5’-CCGGGAGAGT-TTGTGGAAGTCATTTCTCGAGAAATGACTTCCACAAACTCTCTTTTTG-3’ (F), 5’-AAT-TCAAAAAGAGAGTTTGTGGAAGTCATTTCTCGAGAAATGACTTCCACAAACTCTC-3’ (R); shNUDT5-2 – 5’-CCGGATCGTGACAGTCACCATTAACCTCGAGGTTAATGGTGACT-GTCACGATTTTTTG-3’ (F), 5’-AATTCAAAAAATCGTGACAGTCACCATTAACCTCGAG-GTTAATGGTGACTGTCACGAT-3’ (R); shNUDT5-3 – 5’-CCGGCCTACGCTCTAGCACT-GAAACCTCGAGGTTTCAGTGCTAGAGCGTAGGTTTTTG-3’ (F), 5’-AATTCAAAAACC-TACGCTCTAGCACTGAAACCTCGAGGTTTCAGTGCTAGAGCGTAGG-3’ (R); shNUDT9-1 – 5’-CCGGAGGCCCACAGAGGAGCATATACTCGAGTATATGCTCCTCTGTGGGCCTT-TTTTG-3’ (F), 5’-AATTCAAAAAAGGCCCACAGAGGAGCATATACTCGAGTATATGCT-CCTCTGTGGGCCT-3’ (R); shNUDT9-2 – 5’-CCGGCCTCAGATCAGTGAAAGTAATC-TCGAGATTACTTTCACTGATCTGAGGTTTTTG-3’ (F), 5’-AATTCAAAAACCTCAGATC-AGTGAAAGTAATCTCGAGATTACTTTCACTGATCTGAGG-3’ (R). Resulting vectors were transformed into Stbl3 *E. coli* cells, plasmid DNA was purified, and insertions were validated by automated sequencing.

NUDT5 expression plasmids were generated using Gateway Cloning. Briefly, WT NUDT5 (Q9UKK9, aa1-219) was subcloned into pENTR4-N-GFP or pENTR1a-C-GFP entry vectors with SalI/NotI (Thermo Fisher Scientific), as previously (29,65), before validating by automated sequencing. Entry vectors were then transferred to pLenti-CMV-Blast destination vectors (Addgene plasmid # 17451; (66)) using LR Clonase II (Thermo Fisher Scientific) and confirmed by colony PCR.

### Lentiviral production and transduction

To generate stable expression cell lines by lentiviral transduction, lentiviral particles were produced by transfecting subconfluent HEK293T cells with third-generation packaging plasmids (Gag-Pol, Rev, and VSV-G envelope (Addgene plasmids # 12251, 12253, 12259) as before (29) using the calcium phosphate precipitation method (67). Viral supernatants were collected at 48-and 72-hours post-transfection, and target cells were incubated with a 1:1 mixture of lentiviral supernatant and fresh complete growth medium supplemented with 8 μg/mL polybrene. Forty-eight hours post-transduction, cells were re-seeded at low density and subjected to antibiotic selection with either blasticidin (5 μg/mL; Sigma-Aldrich) for four days or puromycin (1 μg/mL; Sigma-Aldrich) for three days, with media replenished every three days.

### Reverse transcription quantitative PCR (RT-qPCR)

Cells were treated as indicated before harvesting with TRIzol (Thermo Fisher Scientific). RNA was purified with the Direct-zol RNA MiniPrep kit (Zymo Research) according to the manufacturer’s instructions and quantified by NanoDrop (Thermo Fisher Scientific). cDNA was then generated with the iScript cDNA Synthesis Kit (Bio-Rad) according to the manufacturer’s instructions. qPCR was performed with 2.5 ng cDNA per sample and iTaq Universal SYBR Green Supermix (Bio-Rad) using a Bio-Rad CFX96 Real-Time PCR Detection System (CFX Maestro™ 1.0 software). Relative quantity of *NUDT5* (F: 5’-TGAAGCTAGGTCAAAGGCGG-3’, R: 5’-GTTCACCTCCAAGGTGTAGGG-3’) and *NUDT9* mRNA (F: 5’-ACCTAGCATCGCAAAG-CCG-3’, R: 5’-ACGAGTTTCTGAACGCCTGG-3’) was calculated using the ΔΔCt method via normalization to *GAPDH* (F: 5’-AAGGTCGGAGTCAACGGATT-3’, R: 5’-CTCCTGGAAG-ATGGTGATGG-3’) and *β-actin* (F: 5’-CCTGGCACCCAGCACAAT-3’, R: 5’-GGGCCGGACT-CGTCATACT-3’).

### Western blotting

Cells were treated as described and harvested in RIPA buffer. Following lysis and clarification, total protein was measured by BCA assay (Pierce), and samples were normalized prior boiling in 1× (v/v final) Laemmli buffer. Protein samples were separated on 4-20% gradient Mini-PROTEAN gels (Bio-Rad) prior to transferring onto 0.2 µm nitrocellulose with a Trans-Blot Turbo Transfer System (Bio-Rad). After blocking with LI-COR Blocking Buffer (TBS; LI-COR) for 1 hour at room temperature, primary antibodies were added at the following concentrations in 1:1 LI-COR Blocking Buffer and TBS + 0.05% Tween-20 at 4°C overnight: anti-GFP (rabbit polyclonal, 1:500), anti-PARP1 (mouse monoclonal, 1:500), anti-PARP1 (rabbit monoclonal, 1:5,000), anti-SOD1 (mouse monoclonal, 1:500), anti-NUDT5 (rabbit polyclonal, 1:1,000), anti-NUDT5 (mouse monoclonal, 1:500), anti-β-actin (mouse monoclonal, 1:5,000), anti-PYCR1 (rabbit polyclonal, 1:2,000), anti-PYCR2 (mouse monoclonal, 1:1,000), and anti-PPAT (rabbit polyclonal, 1:1,000). LI-COR secondary antibodies were diluted in 1:1 LI-COR Blocking Buffer (TBS) and TBS + 0.05% Tween-20 at (1:10,000) prior to incubating at room temperature for 1 hour. Blots were imaged with a LI-COR Odyssey Fc and analyzed using Image Studio (LI-COR, version 5.2).

### Viability assays

#### Resazurin reduction assay

Cells were treated as described in 96- or 384-well back assay plates with clear bottom (BD Falcon or Corning) and viability was measured by resazurin reduction viability assay. For experiments with antifolates, cells were instead grown in complete DMEM high glucose or RPMI media with 10% dialyzed FBS (dFBS, Thermo Fisher Scientific). In general, 1500-4000 cells were plated per well and incubated with test compounds for 96 hours before viability readings. shRNA expression was induced with 0.75 µg/mL doxycycline for 72 hours or NUDT5 compounds were pre-incubated for 24 hours prior to test compound addition. For adherent cells, resazurin stocks were diluted 100-fold in cell culture medium and added to cells after removal of treatment media. For suspension cells, resazurin was diluted 50-fold in culture medium and added 1:1 to treatment medium (96-well plates) or diluted 16.67-fold and added at a factor of 1:5 to treated cultures (384-well plates). The final concentration of resazurin in all cases was 0.01 mg/mL. Following incubation in a humidified incubator for 4-6 hours, resorufin fluorescence was measured on a CLARIOStar microplate reader (BMG LABTECH) with 600±40 nm emission filter or a Hidex Sense microplate reader with 595±10 nm emission filter. Background (medium with resazurin alone) was subtracted from readings prior to normalization to the max viability plateau or DMSO control to yield percent viability. Data were fit to using the [Agonist] vs. response – variable slope (four parameters) curve fitting function in GraphPad Prism (version 10) to calculate EC_50_ values and viability minima (bottom plateau).

#### CellTiter-Glo assay

Cells were treated as indicated above in white, 96-well plates (Greiner) and viability was measured by CellTiter-Glo assay (Promega), according to the manufacturer’s instructions. Bioluminescence was measured on a CLARIOStar microplate reader (BMG LABTECH). Background (medium with CellTiter-Glo reagents alone) was subtracted from readings prior to normalization to the max viability plateau or DMSO control to yield percent viability. Data were fit to using the [Agonist] vs. response – variable slope (four parameters) curve fitting function in GraphPad Prism (version 10) to calculate EC_50_ values and viability minima (bottom).

### Proliferation assays

Cells were grown in complete RPMI GlutaMAX medium without folic acid (FA; Thermo Fisher Scientific) and 10% dialyzed FBS (dFBS; Thermo Fisher Scientific) for 72 hours, where DMSO or NUDT5-targeting molecules were added in the last 24 hours. The cells were then re-seeded into 12-well plates in FA-free RPMI/dFBS alone or supplemented with 2 µmol/L folic acid, 5-formyl-THF, or 5-methyl-THF for an ensuing 72 hours before comparing confluence with a CELLCYTE X (Cytena) live-cell microscope.

### Nucleoside analog library viability screen

#### Screening compound handling and spotting

The Nucleotide Compound Library (HY-L044), consisting of 179 compounds, was purchased from MedChemExpress and housed at the Science for Life Laboratory Compound Center, part of Chemical Biology Consortium Sweden (CBCS). The compounds are kept at -20 °C as solutions in DMSO up to 10 mmol/L and under low humidity using a REMP Small-Size Store system. Phosphorylated analogs were triaged from consideration due to cell uptake concerns (**Supplementary Table S1**), leaving 157 unique molecules (6 duplicates with alternative formulations and 8-chloro-adenosine yields 152 compounds). Stocks were transferred to LabCyte 384 LDV plates (LP-0200) to enable dispensing into assay plates with an Echo 550™ acoustic liquid handler (LabCyte). Compound stock solutions were dispensed into polypropylene V-bottom 384-well plates (Greiner), covered with adhesive foil, and stored at -20°C until use.

#### Screen set-up and execution

HEK293T cells were grown and maintained in DMEM high glucose GlutaMAX with 10% heat-inactivated FBS. On day 0, cells were plated in T25 flasks and pre-treated with DMSO (0.1% v/v), 5 µmol/L TH5427, and 1 µmol/L DDD2 for 24 hours and were maintained in these treatments throughout the experiment. The following day (day 1), 50 µL of cells (500 per well) were added to black, clear-bottomed 384-well cell culture assay plates (BD Falcon) using a VIAFLO 384 (Integra). The compounds were thawed, diluted in 30 µL of culture medium, and transferred to the assay plates using a JANUS 3 automated liquid handling workstation (PerkinElmer) for a final volume of 80 µL per well. Columns 1 and 2 were reserved for DMSO control cells, while column 24 were for no-cell controls. The assay plates with cells and test compounds were then placed in a humidified incubator at 37°C and 5% CO_2_ for 4 days (96 hours). On day 5, 60 µL of culture medium was removed with a BioTek plate washer (Agilent), and 20 µL of 2x resazurin in culture medium was added to each well with a Multidrop Combi reagent dispenser (Thermo Fisher Scientific) for a final volume of 40 µL and 0.01 mg/mL resazurin. The plates were then placed in 37°C humidified incubators for 3 hours before resorufin fluorescence was measured with an EnVision plate reader (PerkinElmer). The average Ź, an indicator of screening assay quality (68), was 0.83 per plate.

### CellMinerCDB data analysis

NCI-60 drug screening datasets pertaining to drug sensitivity (z-score from negative log_10_[GI_50_(molar)] data across NCI-60 for single drugs [sulforhodamine B total protein cytotoxicity assay, 48hrs post-treatment]) as related to protein abundance (SWATH-MS; protein intensity) or mRNA expression (z-score from microarray log_2_ intensity data across NCI-60) for target genes were exported from the CellMinerCDB (version 2.2) (28). Of the 25,293 compounds assayed in the NCI-60 screen, 1,078 were identified as being nucleoside-like (containing purine and/or pyrimidine moieties) and triaged for further analyses (**Supplementary Table S2 and S3**). Pearson correlations (*r*) and linear regressions (± 95% confidence intervals) were computed for individual compounds across the NCI-60 panel but also between gene pairs across the 1,078 nucleoside-like subsets.

### Metabolomics

#### Sample preparation

Prior to quenching, HEK293T cells were treated as indicated and washed three times with 1 mL PBS. Quenching was performed by adding 500 µL 80% methanol containing internal standards (13C3-caffeine, D-sucrose-13C12). Cells were detached from the wells with a cell scraper, mixed, and 400 µL was transferred to Eppendorf tubes. Media blanks were prepared in the same way as the samples. The samples were snap frozen in liquid nitrogen and stored at -80 °C until shipment on dry ice to SMC. Metabolite extraction was performed by adding 100 µL extraction buffer (80:20, v/v methanol:water) containing internal standards (LC-MS: phenylalanine-13C9, cholic acid-D4, caffeic acid-13C9, salicylic acid-D6; GC-MS: L-proline-13C5, alpha-ketoglutarate-13C4, myristic acid-13C3, cholesterol-D7, succinic acid-D4, salicylic acid-D6, L-glutamic acid-13C5,15N, putrescine-D4, hexadecanoic acid-13C4, D-glucose-13C6). The sample was shaken with a tungsten bead at 30 Hz for 3 min in a mixer mill (Retsch GmbH, Haan, Germany), the bead was removed, and the samples were centrifuged at 4 °C, 14,000 rpm (18,620 g) for 10 min (Hettich Mikro 220R). Supernatant fractions (200 µL for LC-MS and 50 µL for GC-MS) were transferred to microvials and evaporated to dryness in a vacuum concentrator (GeneVac miVac Quattro Concentrator, Ipswich, UK). The samples were stored at -80 °C until analysis. Extraction blanks were prepared in the same way as the samples. Small aliquots of the remaining supernatants were pooled to generate QC samples, which were aliquoted and injected at regular intervals to monitor data quality. QC samples were also analysed by LC-MS/MS for metabolite identification. The samples were analysed in batches according to a randomized run order on both GC-MS and LC-MS.

#### LC-MS analysis

Before LC–MS analysis, the sample was re-suspended in 10 + 10 µL methanol and water. Samples were analyzed first in positive ion mode, followed by a second injection in negative ion mode. Chromatographic separation was performed on an Agilent 1290 Infinity UHPLC system (Agilent Technologies, Waldbronn, Germany). 2 µL of each sample was injected onto an Acquity UPLC HSS T3 C18 column (2.1 × 50 mm, 1.8 µm) with a VanGuard precolumn (2.1 × 5 mm, 1.8 µm; Waters Corporation, Milford, MA, USA) held at 40 °C. The gradient solvents were A: H2O with 0.1% formic acid, and B: acetonitrile/2-propanol (75:25, v/v) with 0.1% formic acid. The flow rate was 0.5 mL min–1. Elution was performed with a linear gradient of 0.1–10% B over 2 min, increased to 99% B over 5 min, held for 2 min, reduced to 0.1% B over 0.3 min with a simultaneous increase in flow rate to 0.8 mL min–1 for 0.5 min, held for 0.9 min, and then returned to 0.5 mL min–1 for 0.1 min before the next injection. Detection was carried out on an Agilent 6546 Q–TOF mass spectrometer equipped with a jet stream electrospray ion source operating in positive or negative ion mode. The same settings were used for both modes, except for the capillary voltage. A reference interface was connected for internal mass calibration using purine (4 µM, m/z 121.05 and 119.03632) and HP-0921 (1 µM, m/z 922.0098 and 966.000725) infused at 0.05 mL min–1. The gas temperature was 150 °C, drying gas flow 8 L min–1, and nebulizer pressure 35 psig. The sheath gas temperature was 350 °C with a flow rate of 11 L min–1. The capillary voltage was 4000 V in both modes, the nozzle voltage 300 V, the fragmentor voltage 120 V, the skimmer 65 V, and OCT 1 RF Vpp 750 V. Collision energy was set to 0 V. Data were acquired in centroid mode over m/z 70–1700 at 4 scans s–1 (1977 transients per spectrum).

#### GC-MS analysis

Derivatization and GC–MS analysis were performed as described previously (69). Samples were derivatized in 10 µL methoxyamine (15 µg/µL in pyridine) and 20 µL of a 1:1 mixture of MSTFA (+1% TMCS) and heptane containing methyl stearate (15 ng/µL). 1 µL of the derivatized sample was injected in splitless mode by an L-PAL3 autosampler (CTC Analytics AG, Switzerland) into an Agilent 7890B gas chromatograph equipped with a 10 m × 0.18 mm fused silica capillary column with a chemically bonded 0.18 µm Rxi-5 Sil MS stationary phase (Restek Corporation, USA). The injector temperature was 270 °C, the purge flow rate 20 mL min–1, and the purge was turned on after 60 s. The carrier gas flow rate through the column was 1 mL min–1. The column temperature was held at 70 °C for 2 min, increased by 40 °C min–1 to 320 °C, and held for 2 min. The column effluent was introduced into the ion source of a Pegasus BT time-of-flight mass spectrometer, GC/TOFMS (LECO Corp., St. Joseph, MI, USA). The transfer line and ion source temperatures were 250 °C and 200 °C, respectively. Ions were generated by a 70 eV electron beam at an ionization current of 2.0 mA, and 30 spectra s–1 were recorded over m/z 50–800. The acceleration voltage was turned on after a solvent delay of 150 s, and the detector voltage was 1800–2300 V.

#### Data processing and metabolite annotation

For the GC-MS data, all non-processed MS files from the metabolic analysis were exported from ChromaTOF software in NetCDF format to MATLAB R2021a (MathWorks, Natick, MA, USA), where data pre-treatment procedures such as baseline correction, chromatogram alignment, data compression, and multivariate curve resolution were performed. Extracted mass spectra were identified by comparison of their retention indices and mass spectra with libraries of retention time indices and mass spectra (70). Mass spectra and retention index comparison were performed using NIST MS 2.2 software. Annotation of mass spectra was based on reverse and forward library searches, with particular attention to diagnostic masses and mass ratios indicative of derivatized metabolites. A deconvoluted peak was annotated as a metabolite identification when the best-matching library spectrum had the highest probability and the retention index difference between the sample and library was within ±5 (usually <3).

For the LC-MS data, processing was performed in both targeted and untargeted modes. Targeted analysis was carried out using Agilent MassHunter Profinder version B.10.0.0 (Agilent Technologies Inc., Santa Clara, CA, USA). An in-house LC–MS library, constructed mainly from authentic standards run on the same system with identical chromatographic and mass spectrometric settings, was searched using the Batch Targeted Feature Extraction workflow with an m/z tolerance of 20 ppm and a retention time window of 0.1 min.

For untargeted processing, MZmine 4.4.3 (71) was used. Raw LC-MS data were converted to mzML format using MSConvert (ProteoWizard). Mass detection was performed with an intensity threshold of 2 × 103. The chromatogram builder was set to the Local Minimum Feature Resolver with an m/z tolerance of 20 ppm, peaks were aligned using the Join Aligner algorithm with an m/z tolerance of 20 ppm, and gap filling was performed using the Peak Finder method. A local compound database search was applied to highlight previously annotated metabolites. Group filtering (≥ 50% in one metadata group), signal-to-noise filtering (above 2.5 × extraction blank), QC dilution series correlation filtering (positive correlation), and removal of previously annotated metabolites were performed with an in-house R script (R v.4.4.2, R Core Team, 2025).

Confidence levels for annotation of the metabolites were assigned according to established community systems (72–74). Five levels were defined: Level 1 (confirmed identification), based on direct comparison with an authentic reference standard analyzed under identical conditions, requiring agreement in accurate mass, retention time, and usually MS/MS spectra; Level 1* (isomeric mixture), applied when standards confirmed a mixture of structural isomers without chromatographic or MS/MS separation; Level 2 (putative annotation/structure), based on spectral library matches and/or diagnostic MS/MS evidence but without a standard; Level 3 (tentative annotation/compound class), based on accurate mass, isotope distribution, and database search, enabling tentative structure or class assignment; Level 4 (molecular formula), features with accurate mass and molecular formula but insufficient evidence for structure; and Level 5 (unique feature), accurate mass only, without confirmed formula.

### Proteomics

#### Sample processing

Proteomic analysis was performed to assess compound-induced changes in total protein abundance but also changes in soluble protein abundance by Proteome Integral Solubility Alteration (PISA) assay, using tandem mass tag (TMTpro)-based quantitative mass spectrometry, as previously described (75–77). In this assay, named as “PISA-Express”, each biological replicate was treated either with DDD2 or DMSO in culture and halved for each analysis as an independent sample in the same TMT-based multiplex: one half is subjected to thermal treatment and protein extraction for PISA assay, while the other is processed equally but for total protein expression (and normalization of soluble protein by PISA). HEK293T cells were treated with either vehicle (DMSO, 0.1% v/v) or 1 µmol/L DDD2 – with four experimental replicates per condition – for 24 hours, detached and resuspended in PBS with protease inhibitors before processing, as above. After protein extraction by freeze-thaw cycles in liquid nitrogen and 37°C with 0.4% NP-40, the soluble fraction was collected by ultracentrifugation and quantified by microBCA assay (Thermo Fisher). Samples were then reduced, alkylated, acetone-precipitated, and digested with LysC and trypsin. Each digest was labeled using TMTpro technology (Thermo Fischer) for deep protein identification and quantification by LC-MS data-dependent acquisition on a 16-plexed sample constituted of 4 replicates of DMSO and 4 replicates of DDD2-treated samples for both PISA assay and expression proteomics. The final multiplexed sample was first cleaned by Sep-Pack C18 column (Waters) and then separated into 48 fractions by capillary reversed-phase chromatography at pH 10. Each fraction was analyzed by high-resolution nLC–ESI-MS/MS (nanoscale liquid chromatography-electrospray ionization-tandem mass spectrometry) using an Orbitrap Exploris 480 instrument (Thermo Scientific) (76,78).

#### NanoLC-MS/MS analysis

NanoLC-MS/MS analyses were performed on an Orbitrap Exploris 480 mass spectrometer (Thermo Fisher Scientific). The instrument was equipped with an EASY ElectroSpray source and connected online to an Ultimate 3000 nanoflow UPLC system. The samples were pre-concentrated and desalted online using a PepMap C18 nano-trap column (length - 2 cm; inner diameter -75 µm; particle size -3 µm; pore size -100 Å; Thermo Fisher Scientific) with a flow rate of 3 µL/min for 5 min. Peptide separation was performed on an EASY-Spray C18 reversed-phase nano-LC column (Acclaim PepMap RSLC; length -50 cm; inner diameter -2 µm; particle size - 2 µm; pore size – 100 Å; Thermo Scientific) at 55 °C and a flow rate of 300 nL/min. Peptides were separated using a binary solvent system consisting of 0.1% (v/v) FA, 2% (v/v) ACN (solvent A) and 98% ACN (v/v), 0.1% (v/v) FA (solvent B). They were eluted with a gradient of 3-26% B in 97 min, and 26-95% B in 9 min. Subsequently, the analytical column was washed with 95% B for 5 min before re-equilibration with 3% B. The mass spectrometer was operated in a data-dependent acquisition mode. A survey mass spectrum (from m/z 375 to 1500) was acquired in the Orbitrap analyzer at a nominal resolution of 120,000. The automatic gain control (AGC) target was set as 100% standard, with the maximum injection time of 50 ms. The most abundant ions in charge states 2+ to 7+ were isolated in a 3 s cycle, fragmented using HCD MS/MS with 33% normalized collision energy, and detected in the Orbitrap analyzer at a nominal mass resolution of 50,000. The AGC target for MS/MS was set as 250% standard with a maximum injection time of 100 ms, whereas dynamic exclusion was set to 45 s with a 10-ppm mass window.

#### Protein Identification, Quantification, and Data Analysis

Proteome Discoverer 2.5 software (Thermo Scientific) was utilized for protein database search, peptide/protein identification and quantification against the Uniprot Homo sapiens (Human) protein database (UP000005640). Cysteine carbamidomethylation was set as a fixed modification, along with TMT-related modifications, methionine oxidation, deamidation of arginine and asparagine as variable modifications. Enzyme specificity was defined as trypsin with a maximum of two missed cleavages. A 1% false discovery rate was employed as a filter at both the protein and peptide levels. Contaminants and reversed-hit peptides were removed, and only proteins with at least two unique peptides were included in the quantitative analysis. Proteins with missing values were also eliminated. The quantified abundance of each protein in each sample (labeled with a different TMTpro reagent) was normalized to the total intensity of all proteins in that sample. For each protein in each treatment replicate, the normalized protein abundance was divided by the average abundance of that protein in the vehicle-treated replicates. Soluble protein abundance in the PISA assay was calculated by normalizing to the total amount from the respective expression proteomics. The average ratio across replicates of each compound compared to the vehicle control was calculated, and the Log_2_ values of these ratios were determined. Two-tailed Student’s t-test was employed to calculate the p-value, assuming a non-zero ratio. For each compound, the results were visualized using a volcano plot, which plotted the Log_2_ ratio on the x-axis and the corresponding -Log_10_ p-value on the y-axis. The mass spectrometry proteomics data have been deposited to the ProteomeXchange Consortium *via* the PRIDE partner repository (79) with the dataset identifier PXD069745.

### Co-immunoprecipitation

#### GFP-Trap

Co-immunoprecipitation of GFP fusion protein interactors from C-GFP, GFP-NUDT5, and NUDT5-GFP-expressing HEK293T cells was performed with GFP-Trap Magnetic Agarose Kit (ChromoTek) according to the manufacturer’s instructions. Portions of the input lysate (IN), immunoprecipitated complexes (IP), and unbound eluate (flowthrough, FT) were reserved for western blotting analysis after addition of 2x Laemmli buffer and heating at 95°C for 5 minutes.

#### Endogenous NUDT5 complexes

Co-immunoprecipitation of endogenous NUDT5 complexes was performed with Protein G Magnetic Beads (Thermo Fisher Scientific), according to the manufacturer’s instructions. Briefly, ∼1×10^7^ HL-60 cells were lysed per immunoprecipitation reaction in ice-cold 1% NP-40 buffer (50 mmol/L Tris-HCl pH 8.0, 150 mmol/L NaCl, 1% NP-40, cOmplete EDTA-free protease inhibitor [Roche], PhosSTOP phosphatase inhibitor [Roche]) for 30 minutes on ice. Samples were clarified by centrifugation for 20 minutes at 20,000 x *g* and 4°C before quantitation by BCA assay (Pierce). Then, 1 mg protein was incubated with 7.5 µg of NUDT5 antibody (rabbit polyclonal, rabbit polyclonal [Abcam], or mouse monoclonal [Santa Cruz]) overnight at 4°C with gentle mixing. The following morning, 25 µL Protein G Magnetic Beads (∼0.25 mg; Pierce) were washed with TBS + 0.05% Tween-20 and 0.5 M NaCl before addition of samples and gentle mixing for 1 hour at room temperature. Portions of the input lysate, immunoprecipitates, and flow-through were collected and prepared for western blotting as above.

### AlphaFold Server complex modeling and structure visualization

Protein complexes were modeled using AlphaFold Server (AlphaFold 3) (48). Input wild-type sequences were provided for PPAT (Q06203, 517 AAs) and NUDT5 (Q9UKK9, 219 AAs) and modeled in different multimeric states. For the effects of NUDT5 Tyr74 phosphorylation, complexes were modeled based on NUDT5 with a phosphorylation at Tyr74. For predicted structures, pTM (accuracy of the overall structure), ipTM (precision of individual components within the entire structure), and PAE (pairwise confidence of amino acid positioning within the structure) are provided to indicate the overall confidence of predicted structures. For structures with poorer overall confidence scores (*ie*, NUDT5 complexes with active PPAT dimers), pIDDT (per-atom confidence score) is also provided.

Structures for *E. coli* (PDBID: 1ECF) (50) and *B. subtilis* (PDBID: 1AO0) (49) were visualized and prepared for graphics using Protein Imager (80).

### Statistical analysis

GraphPad Prism was used for statistical analyses. Specifics on statistical testing for different experiments can be found in the respective methodological sections or in the figure legends.

## Acknowledgements

We would like to express our gratitude to the SciLifeLab Compound Center, part of the Chemical Biology Consortium Sweden (CBCS), for assistance with compound management and spotting. Swedish Metabolomics Centre, Umeå, Sweden (www.swedishmetabolomicscentre.se) is acknowledged for metabolic profiling by GC-MS and LC-MS. Chemical Proteomics at the Karolinska Institutet (Chemistry I Division, MBB Department), also Unit of SciLifeLab and node of the Swedish National Infrastructure for Biological Mass Spectrometry (BioMS), provided full support in the experimental design, performance, and data analysis of the proteomic studies.

## Author contributions

Conceptualization (NCKV, MA), Methodology (NCKV, SA, FMH, UM, JB), Validation (NCKV, SA, FMH, MG), Formal analysis (NCKV, SA, FMH, UM, MG), Investigation (NCKV, SA, FMH, UM, MG), Resources (NCKV, MA, PIA, SGR, BL, MG), Writing – original draft (NCKV), Writing – review & editing (All authors), Visualization (NCKV, SA), Supervision (NCKV, MA, PIA, SGR, BL), Project administration (NCKV, BL, MG, MA), Funding acquisition (NCKV, MA, PIA, SGR).

## Data availability

The proteomics data generated in this study are publicly available in PRIDE at https://www.ebi.ac.uk/pride/markdownpage/proteomexchange, in addition to Supplementary Tables S5 and S6, while metabolomics datasets are available in Supplementary Table S4. Other data are available in the article and its supplementary data files or is available upon request from the corresponding authors.

