## Supplementary Figures for "NUDT5 regulates the global efficacy of nucleoside analog drugs by coordinating purine synthesis and PRPP allocation"

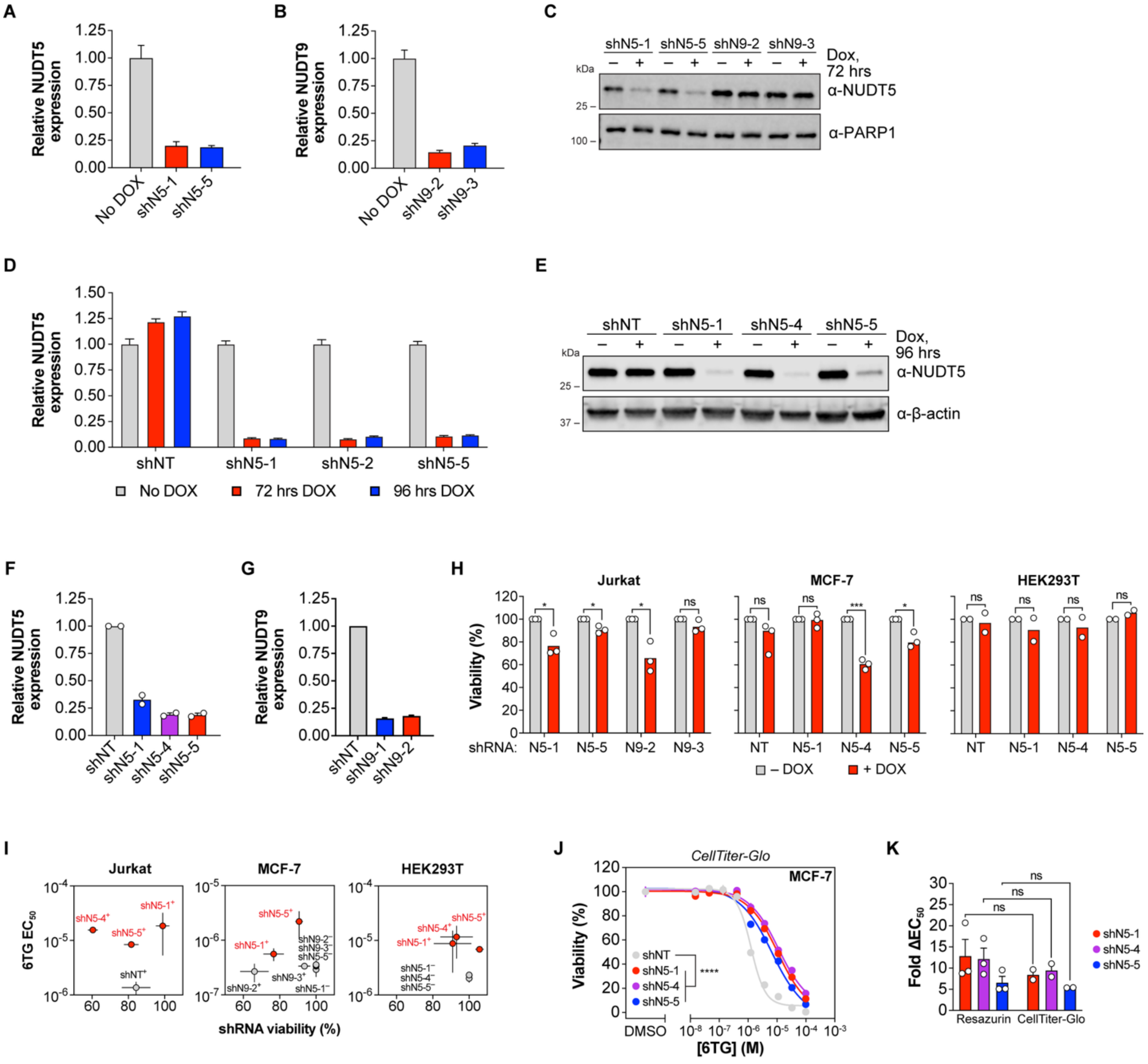


**Supplementary Fig. S1. NUDT5 and NUDT9 shRNA characterization. A**, Relative NUDT5 and NUDT9 (**B**) expression in Jurkat cells following relevant shRNA induction for 72 hours by RT-qPCR. Mean values from a representative experiment performed in triplicate is shown ± SEM. Expression normalized to ß-actin and set relative to “No DOX” control. **C**, NUDT5 protein expression in Jurkat cells following shRNA induction for 72 hours by western blot. A representative blot from n=2 is shown. **D**, Relative NUDT5 expression following shRNA induction in MCF-7 cells for 72 or 96 hours by RT-qPCR. Mean values from a representative experiment performed in triplicate is shown ± SEM. Expression normalized to ß-actin and set relative to “shNT” control. **E**, NUDT5 protein expression in MCF-7 cells following shRNA induction for 96 hours by western blot. A representative blot from n=2 is shown. **F**, Relative NUDT5 and NUDT9 (**G**) expression following relevant shRNA induction in HEK293T cells for 72 hours by RT-qPCR. Mean values from n=2 experiments (NUDT5) or a representative experiment (NUDT9) performed in triplicate is shown ± SEM. Expression normalized to ß-actin and set relative to “shNT” control.

**H**, Cell viability relative to "No DOX" control after 72 hours. Means of n=3 (Jurkat, MCF-7) or n=2 (HEK293T) experiments shown. **I**, shRNA viability *vs* 6TG $EC_{50}$ plots. Means of n=3 (Jurkat, MCF-7) or n=2 (HEK293T) experiments ± SEM are shown. NUDT5 shRNAs are highlighted in red. **J**, NUDT5 was depleted by doxycycline (DOX)-inducible shRNA in MCF-7 cells (n=2) and incubated with a 6TG concentration gradient for 96 hours before viability measurement by CellTiter-Glo. **K**, Comparison of 6TG $EC_{50}$ fold change following NUDT5 shRNA between resazurin (n=3) and CellTiter-Glo (n=2). Statistical significance determined by multiple unpaired, two-tailed t test (**H**, **K**) or extra sum-of-squares F-test for $EC_{50}$ difference from control (**J**), and all errors shown as ± SEM.

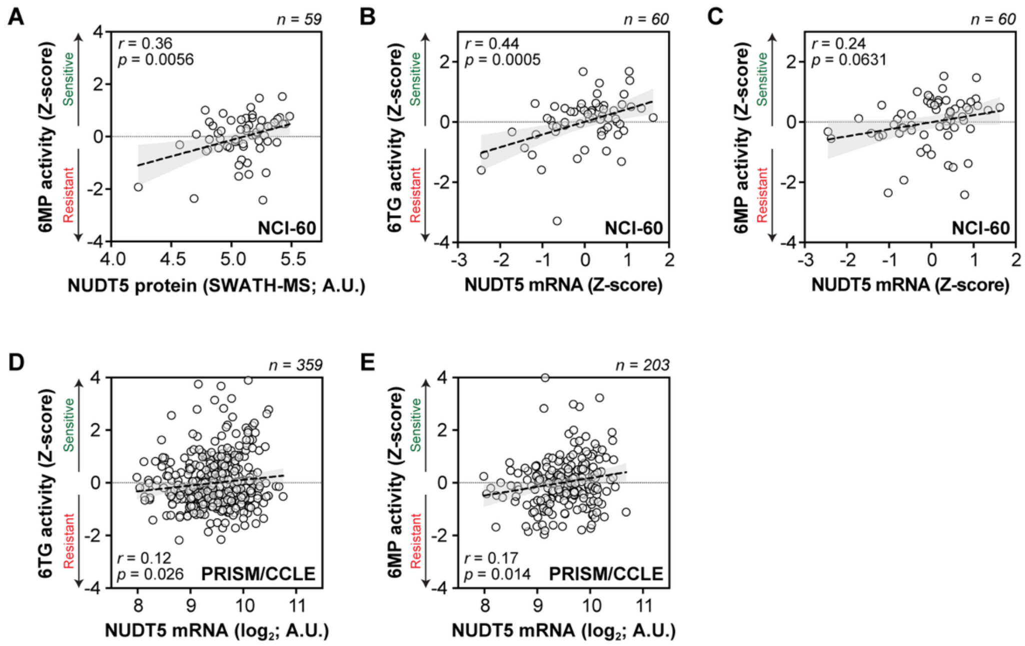


**Supplementary Fig. S2. Correlation of thiopurine activity with NUDT5 expression in CellMinerCDB database. A**, 6MP *vs* NUDT5 protein expression (SWATH-MS) in the NCI-60 dataset (n=59 cell lines). **B**, 6TG *vs* NUDT5 mRNA expression (Z-score) in the NCI-60 dataset (n=60 cell lines). **C**, 6MP *vs* NUDT5 mRNA expression (Z-score) in the NCI-60 dataset (n=60 cell lines). **D**, 6TG *vs* NUDT5 mRNA expression ($log_2$) in the PRISM/CCLE dataset (n=359 cell lines). **E**, 6MP *vs* NUDT5 mRNA expression ($log_2$) in the PRISM/CCLE dataset (n=203 cell lines). For drug sensitivity Z-scores, positive values are considered "sensitive" and negative values "resistant". Pearson correlations (*r*) and associated p-values (two-sided t test) with linear regression ± 95% CI provided for each comparison.

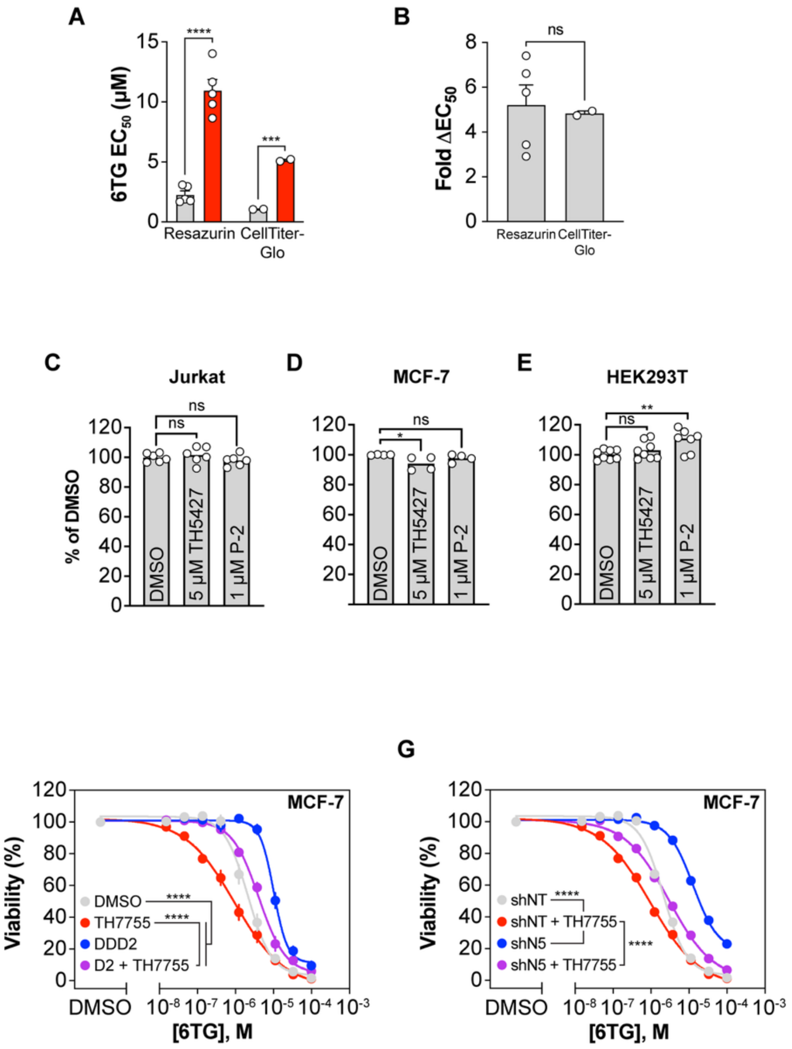


**Supplementary Fig. S3. 6TG resistance in NUDT5-depleted cells is independent of cell viability and NUDT15.** **A**, Comparison 6TG $EC_{50}$ values in DMSO- (grey) or DDD2-treated cells (red) by resazurin reduction (n=5) or CellTiter-Glo assay (n=2). **B**, 6TG $EC_{50}$ fold change between resazurin (n=5) and CellTiter-Glo (n=2) assays. **C**, Viability of Jurkat (n=6), MCF-7 (**D**, n=4), HEK293T (**E**, n=7) cells incubated with DMSO, TH5427, or DDD2 for 96 hours by resazurin assay. **F**, MCF-7 cell viability following DMSO, TH7755, DDD2, or DDD2 + TH7755 for 24 hours and a 6TG gradient for an additional 96 hours (n=3). **G**, MCF-7 cell viability following induction of shNT or shNUDT5-1 for 72 hours and a 6TG gradient ± TH7755 for an additional 96 hours (n=3). Statistical significance determined by multiple unpaired, two-tailed t test (**A**), two-tailed t test (**B**), one-way ANOVA with multiple comparisons to the DMSO control (**C**, **D**, **E**), or extra sum-of-squares F-test for $EC_{50}$ difference from DMSO or shNT control (**F**, **G**), and all errors shown as ± SEM.

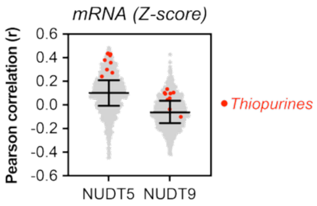


**Supplementary Fig. S4. NA drug efficacy correlations with NUDT5 and NUDT9 expression in the NCI-60.** Pearson correlations (*r*) of 1,078 NA-like drug activities with NUDT5 or NUDT9 mRNA expression (Z-score) in the NCI-60 panel. Thiopurines are highlighted in red, while bars denote median values ± interquartile range.

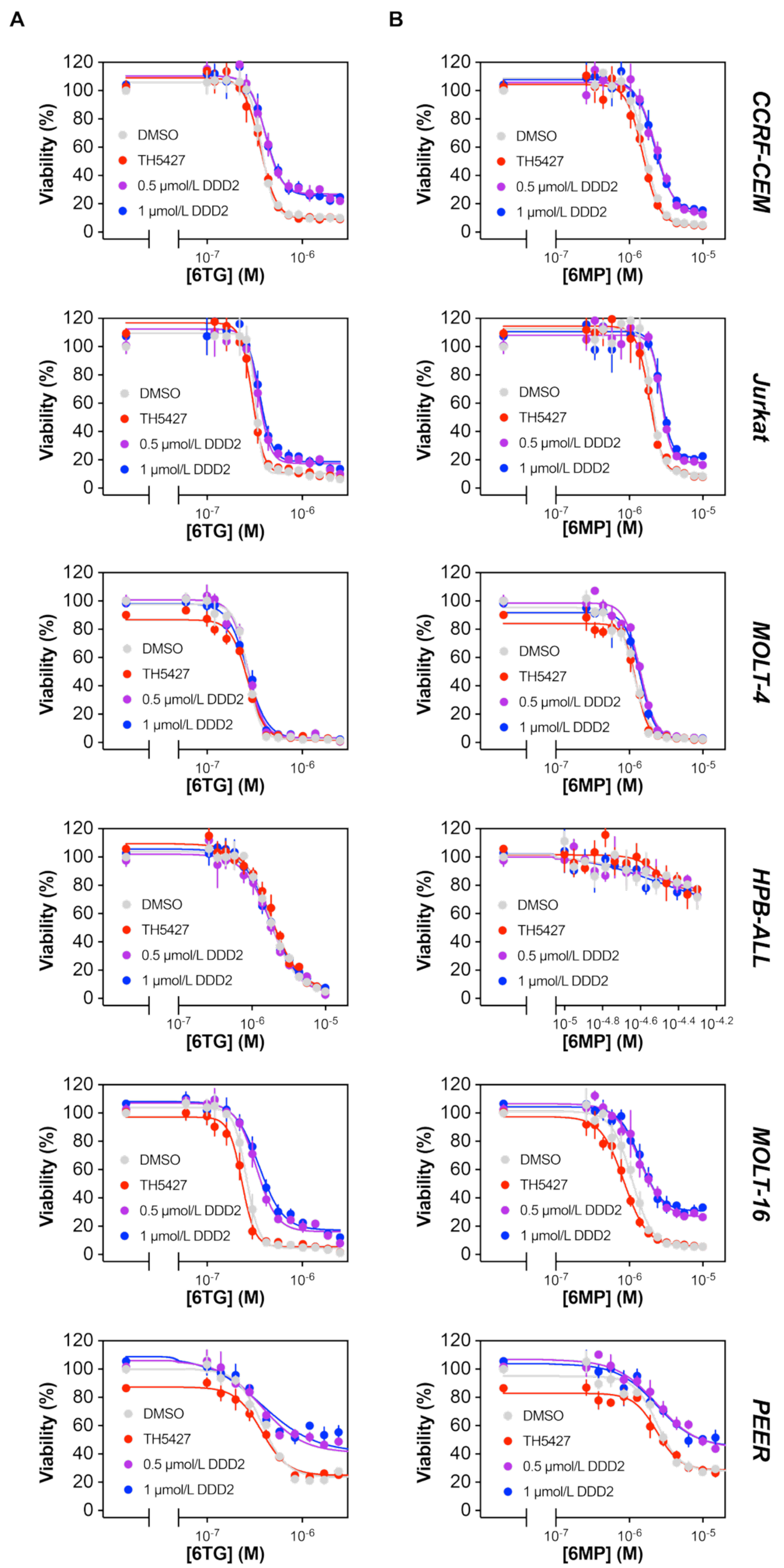


**Supplementary Fig. S5. Effects of NUDT5 inhibition *vs* depletion on thiopurine efficacy in ALL cell lines. A**, CCRF-CEM, Jurkat, MOLT-4, HPB-ALL, MOLT-16, and PEER cells were pre-treated with DMSO, 5 µmol/L TH5427, or 0.5/1 µmol/L DDD2 for 24 hours prior to addition of 6TG or 6-mercaptopurine (6MP; **B**) gradients for an additional 96 hours. Viability was then determined by resazurin reduction assay with normalization to DMSO control values. Means ± SEM with curve fittings are shown from n=3 independent experiments.

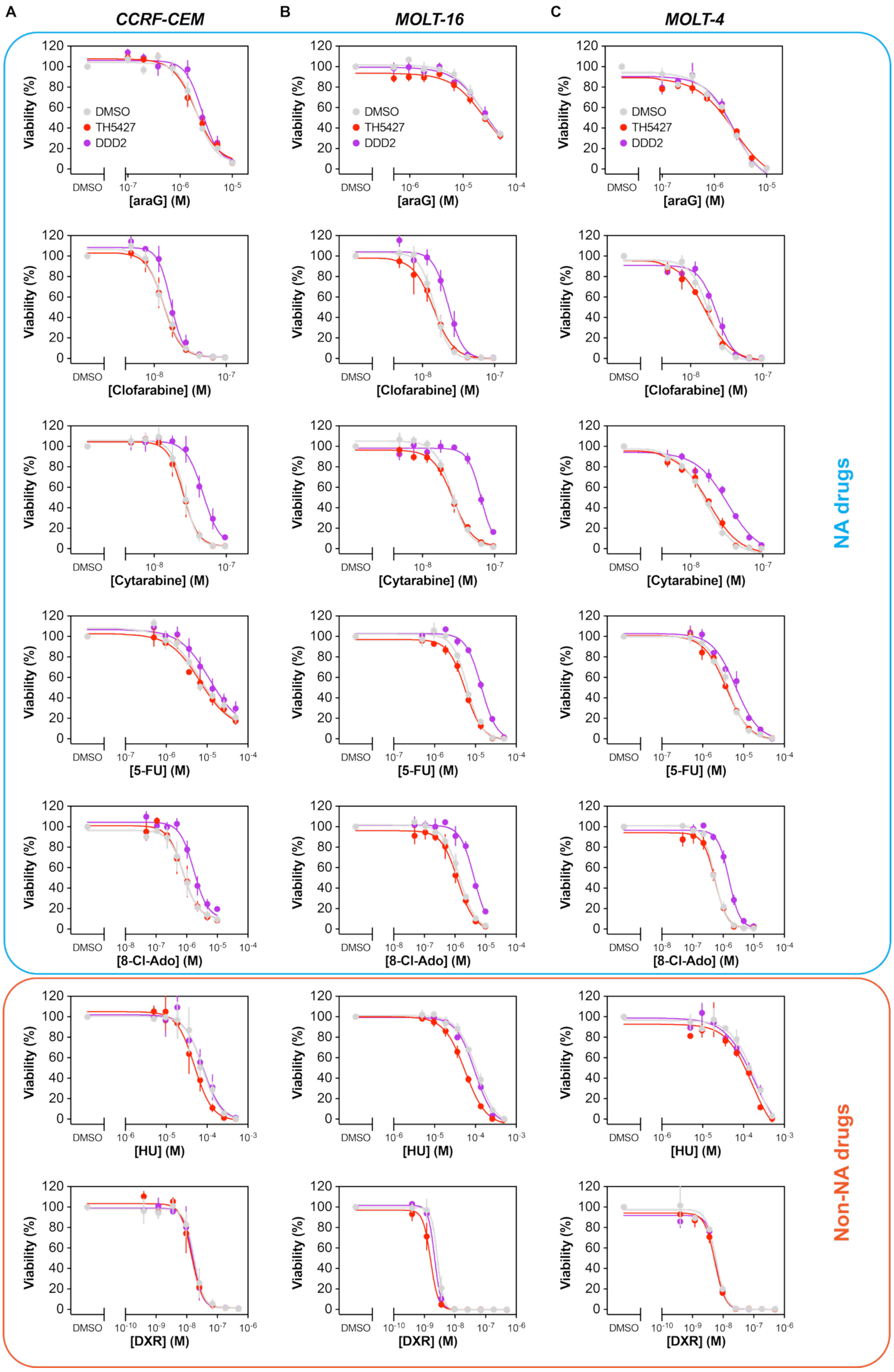

A
CCRF-CEM
B
MOLT-16
C
MOLT-4
DMSO
TH5427
DDD2
Viability (%)
[araG] (M)
[Clofarabine] (M)
[Cytarabine] (M)
[5-FU] (M)
[8-Cl-Ado] (M)
[HU] (M)
[DXR] (M)
NA drugs
Non-NA drugs

**Supplementary Fig. S6. Effects of NUDT5 inhibition *vs* depletion on non-thiopurine NA drug efficacy in ALL cell lines.** **A**, CCRF-CEM, MOLT-16 (**B**), or MOLT-4 (**C**) cells were pre-treated with DMSO, 5 µmol/L TH5427, or 0.5 µmol/L DDD2 for 24 hours prior to addition of NA drug (ara-G, clofarabine, cytarabine [ara-C], 5-fluorouracil (5-FU), 8-chloroadenosine [8-Cl-Ado]) or non-NA cytotoxic drugs (hydroxyurea [HU], doxorubicin [DXR]) for an additional 96 hours. Viability was then determined by resazurin reduction assay with normalization to DMSO control values. Means ± SEM with curve fittings are shown from n=3 independent experiments.

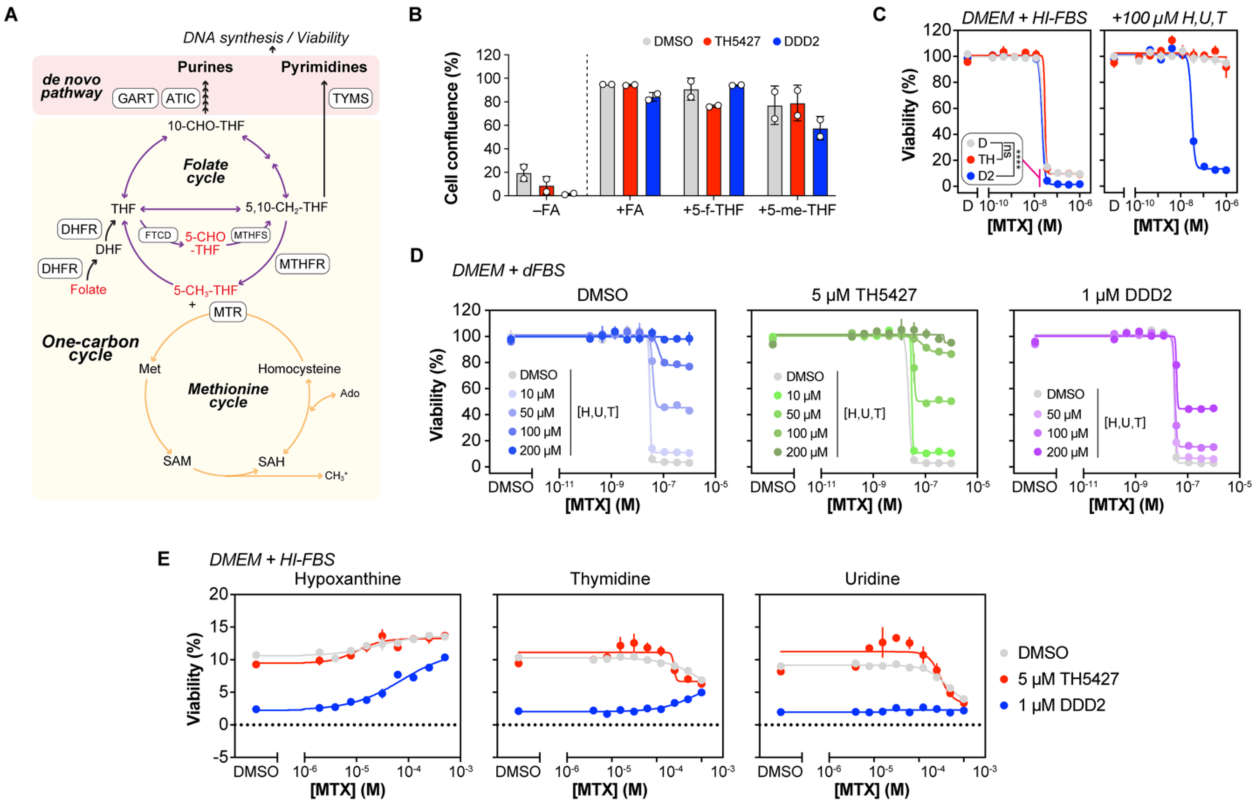


**Supplementary Fig. S7. Rescue of folate deprivation in NUDT5-inhibited or -depleted cells.** **A**, Intersection of one-carbon cycle with *de novo* nucleotide metabolism to support DNA synthesis and proliferation with folate metabolites used for rescue experiments highlighted in red. **B**, DMSO, TH5427, or DDD-2 treated HEK293T cells were grown in folate-free RPMI or folate-free RPMI supplemented with folic acid, 5-formyl-THF, or 5-methyl-THF for 72 hours prior to cell confluence measurements (n=2). **C**, HEK293T cells grown in DMEM with 10% HI-FBS were pre-incubated with DMSO, TH5427, or DDD2 for 24 hours prior to methotrexate (MTX) gradients ± 100 μmol/L hypoxanthine, uridine, thymidine (H, U, T) for an additional 96 hours before resazurin viability measurement. **D**, HEK293T cells grown in DMEM with 10% dFBS were pre-incubated with DMSO, TH5427, or DDD2 for 24 hours prior to methotrexate (MTX) gradients ± H, U, T gradients up to 200 μmol/L for an additional 96 hours before resazurin viability measurement. **E**, HEK293T cells grown in DMEM with 10% HI-FBS were pre-incubated with DMSO, TH5427, or DDD2 for 24 hours prior to methotrexate (MTX) gradients ± individual H, U, or T gradients for an additional 96 hours before resazurin viability measurement. In all cases, statistical significance determined by extra sum-of-squares F-test for bottom difference compared to DMSO control (**C**). Means ± SEM are shown in **B**-**E**.

**A**

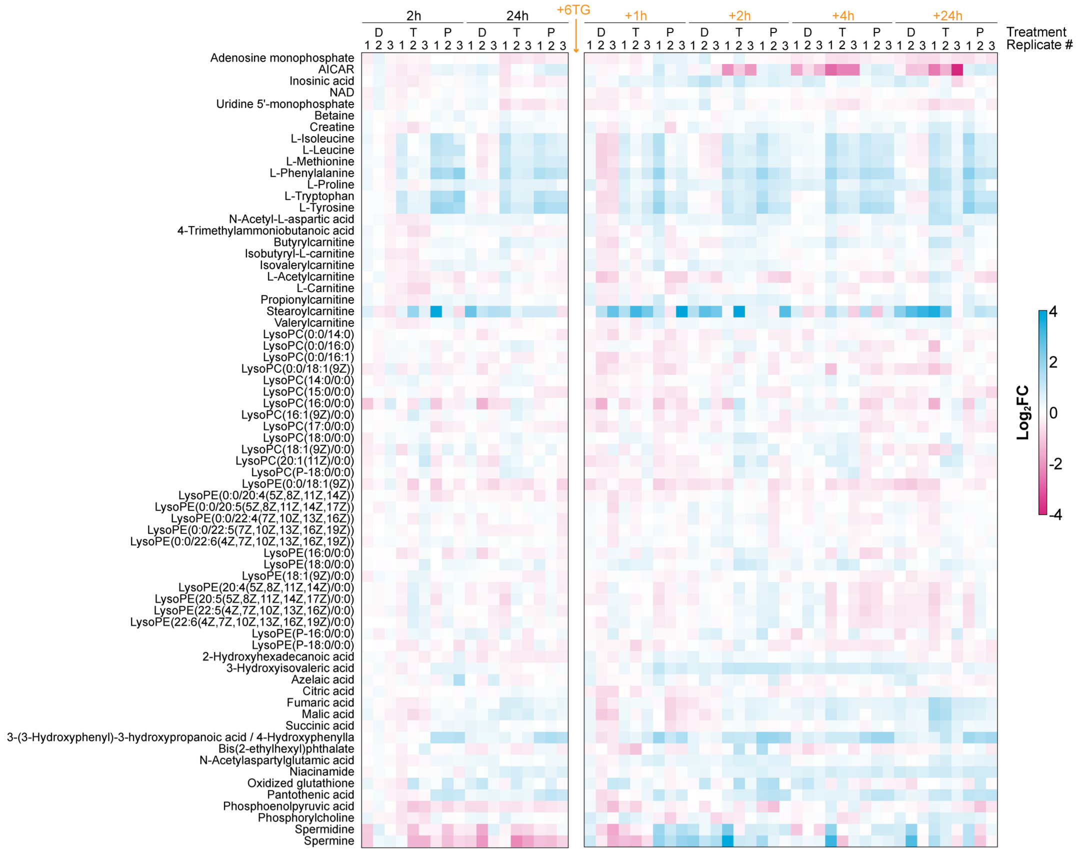


**B**

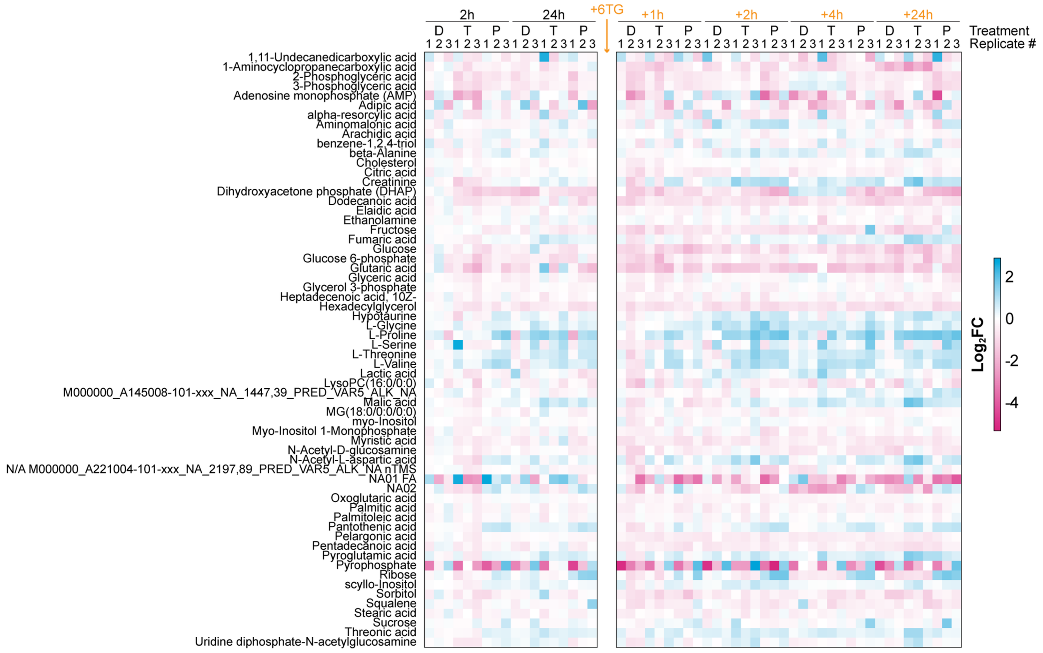


**Supplementary Fig. S8. Metabolomics supplement. A**, Individual replicates (n=3) from LC/MS metabolomics analyses with all metabolites listed. **B**, Individual replicates (n=3) from GC/MS metabolomics analyses with all metabolites listed.

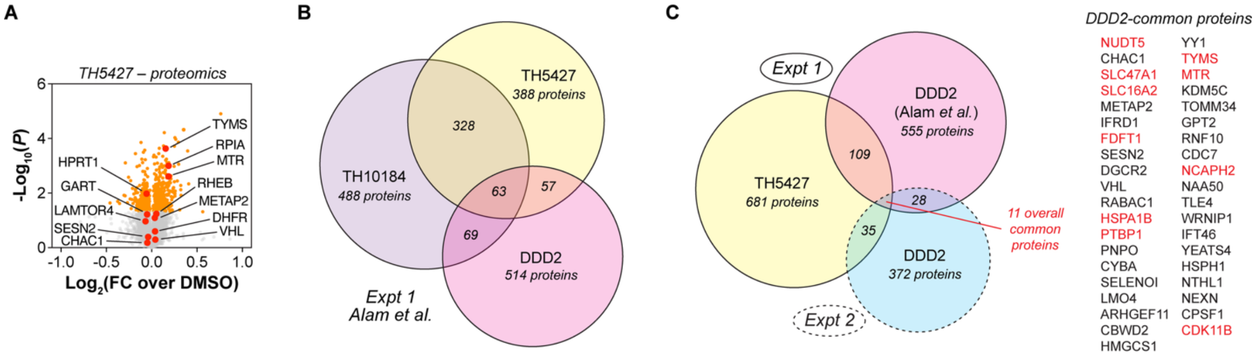


**Supplementary Fig. S9. Comparative proteomics with NUDT5i. A**, Global proteomics volcano plot of protein abundance changes after 5 µmol/L TH5427 for 24 hours. Significant hits shown in orange and highlighted hits in red. Means from n=3 experiments plotted. **B**, Venn diagram of abundance change hits from DDD2- or TH10184-treated cells (Alam *et al.*) and TH5427-treated cells in proteomics experiment 1 with number of proteins labeled in each section. Common proteins are shown from n=3 replicates. **C**, Venn diagram of hits from DDD2- (Alam *et al.*), TH5427-treated cells from experiment 1 and DDD2-treated cells from experiment 2 with number of proteins labeled in each section. Common proteins are shown from n=3 or 4 replicates, with DDD2-common proteins listed and overall common proteins highlighted in red.

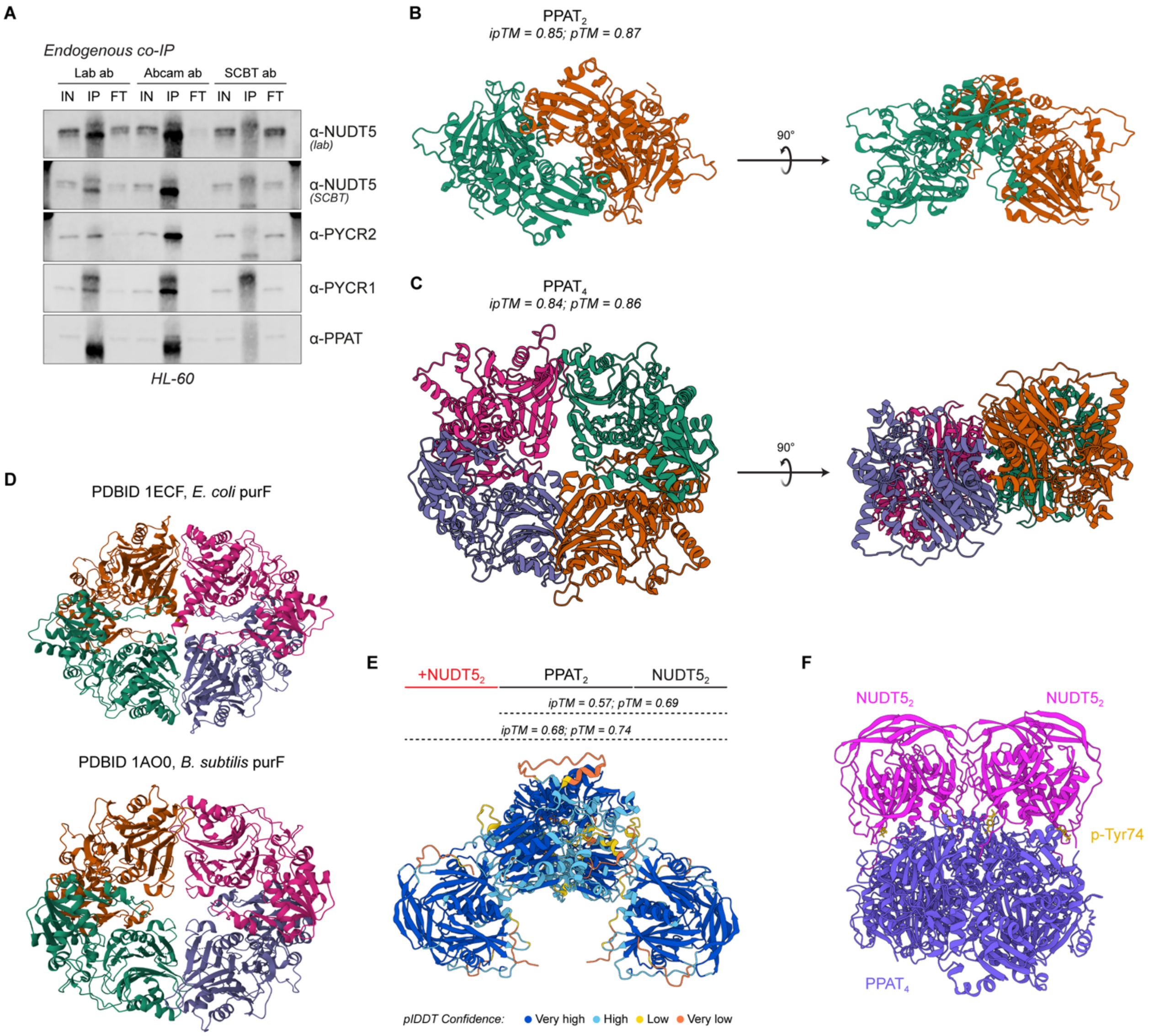


**Supplementary Fig. S10. Modeling of PPAT and PPAT-NUDT5 complexes with AlphaFold 3.** **A**, Co-immunoprecipitation of endogenous NUDT5 with PYCR1/2 and PPAT by three different NUDT5 antibodies. IN – input, IP – immunoprecipitation, FT – flow-through. **B**, High confidence PPAT dimer modeling with AlphaFold 3. **C**, High confidence PPAT tetramer modeling with AlphaFold 3. **D**, Reported X-ray crystal structure of the *E. coli* and *B. subtilis* purF tetramer (PDBID 1ECF and 1AO0, respectively) as visualized on the RCSB webserver. **E**, AlphaFold 3 modeling of potential PPAT-NUDT5 complexes with PPAT in its dimeric configuration and associated prediction confidence scores. **F**, AlphaFold 3 modeling of a potential $PPAT_4$•$(NUDT5_2)_2$ complex in purple and magenta, respectively, with NUDT5 phosphorylated at Tyr74 (yellow).

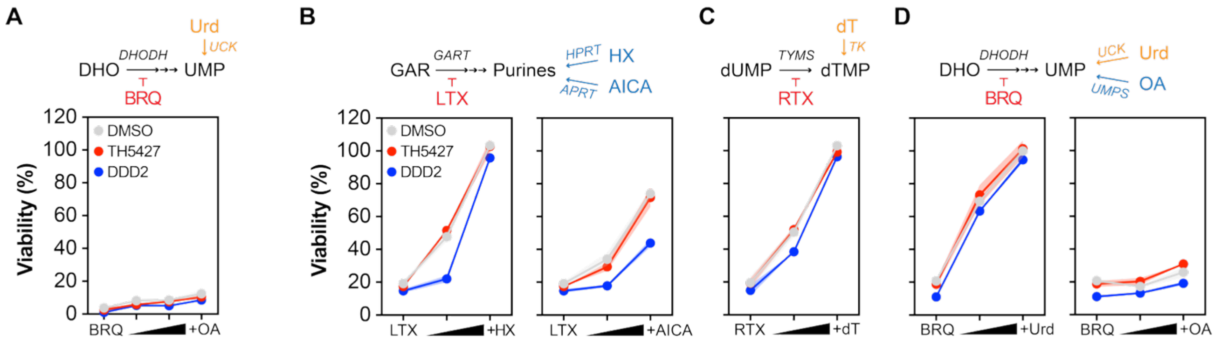


**Supplementary Fig. S11. Purine NA drug sensitivity in control or DDD2-treated in complete or dialyzed FBS. A**, HEK293T cells were grown in DMEM with dialyzed FBS before pre-treatment with DMSO (1% v/v final; grey), 5 µmol/L TH5427 (red), or 1 µmol/L DDD2 (blue) for 24 hours, followed by 1.11 µmol/L brequinar (BRQ) ± 0.5, 0.75, or 1 mmol/L orotic acid (OA) for an additional 96 hours. Viability was then determined by resazurin reduction assay. **B**, Cells were treated as in **A** but with 111 nmol/L lometrexol (LTX) ± 10/100 µmol/L hypoxanthine (HX) or ± 10/100 µmol/L 5-amino-3H-imidazole-4-carboxamide (AICA). **C**, Cells were treated as in **A** but with 1.11 µmol/L raltitrexed (RTX) ± 1/10 µmol/L thymidine (dT). **D**, Cells were treated as in **A** but with 1.11 µmol/L BRQ ± 10/100 µmol/L uridine (Urd) or ± 0.5/1 mmol/L OA. Data shown as means ± SEM from n=2-3 independent experiments.

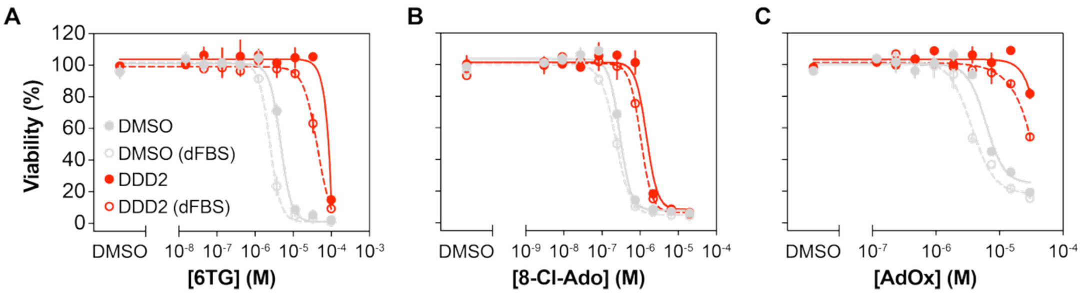


**Supplementary Fig. S12. Purine NA drug sensitivity in control or DDD2-treated in complete or dialyzed FBS. A**, HEK293T cells were grown in DMEM with heat-inactivated or dialyzed FBS before pre-treatment with DMSO (1% v/v final) or 1 µmol/L DDD2 for 24 hours, followed by a 6TG, 8-chloroadenosine (8-Cl-Ado; **B**), or adenosine dialdehyde (AdOx; **C**) concentration gradient for an additional 96 hours. Viability was then determined by resazurin reduction assay.

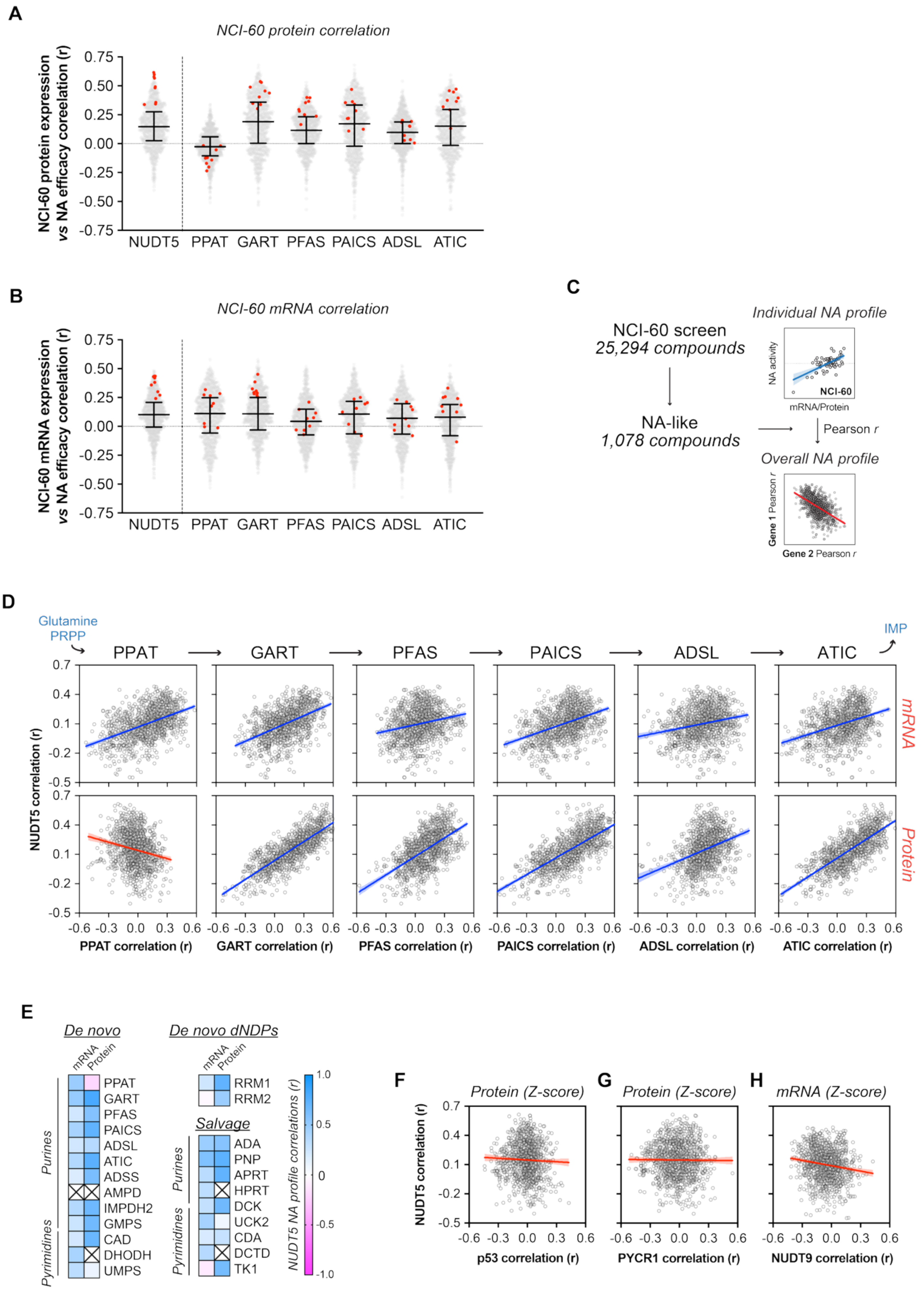


**Supplementary Fig. S13. NA-like drug activity correlations with DNPB enzymes and non-nucleotide metabolizing proteins. A**, Activity correlations of 1,078 NA-like drugs with protein or mRNA (**B**) expression levels for NUDT5 and DNPB enzymes across the NCI-60 panel. Thiopurines are highlighted in red, while bars denote median values ± interquartile range. **C**, Activity-expression correlations of triaged NA-like compounds from the NCI library to determine NA metabolic similarity between two genes. **D**, Correlation of NUDT5-dependent and DNPB enzyme-dependent efficacy correlations of 1,078 NA-like drugs across the NCi-60 cell line panel based on mRNA (top) or protein (bottom) expression from CellMinerCDB with linear regressions ± 95% CI shown. **E**, Pearson correlation heatmap of NUDT5-dependent NA drug efficacy *vs* NA drug efficacy of individual enzymes in nucleotide metabolism based on mRNA or protein abundance to indicate metabolic similarity. Crossed out boxes indicate at least one dataset missing. **F**, Per-compound comparison of NUDT5 with p53, PYCR1 (**G**), or NUDT9 (**H**) NA drug activity Pearson correlations (*r*) based on protein expression for 1,078 NA-like compounds in the NCI-60 panel with linear regression ± 95% CI shown.
